# Dengue virus-induced thrombocytopenia is mediated by dysregulation of the RUNX1–MYH9 axis

**DOI:** 10.64898/2026.09.09.750054

**Authors:** Anjali Singh, Simran Mehra, Kinjal Singh, Sushant Phadnis, Akhil Kumar, Vineet Choudhary, Sankar Bhattacharyya, Bhupendra Verma

## Abstract

Dengue virus (DENV) infection causes an acute febrile illness that is usually self-limiting but can become fatal. Thrombocytopenia, or a decrease in circulating platelets, is a major clinical feature of dengue and is strongly associated with the progression to dengue hemorrhagic fever and dengue shock syndrome. In this report, MYH9, a cytoskeletal protein involved in platelet maturation and release, was found to be down-regulated in DENV-infected *Huh7* cell lines. The expression of RUNX1, an important transcription factor involved in megakaryopoiesis, was also decreased following DENV-infection. *In-silico* analysis identified three potential RUNX1 transcription-factor binding sites in the promoter region of the MYH9 gene. We therefore investigated whether RUNX1 regulates MYH9 expression during DENV-infection. Luciferase and ChIP assays showed that DENV infection affects RUNX1 binding to the MYH9 promoter, providing insight into the molecular basis of reduced MYH9 expression. Furthermore, the megakaryocyte differentiation markers GATA1, ITGB3, and MYH9 were upregulated in PMA-induced K562-cells. However, DENV infection downregulated these genes and increased the expression of the apoptosis marker gene CYCS. These findings suggest that dengue affects megakaryopoiesis through multiple mechanisms, potentially reducing platelet production while also promoting platelet destruction. Additionally, Fluorescence and electron microscopy revealed that functional defects of MYH9 may affect megakaryopoiesis biogenesis. Overall, this study provides insights into the molecular mechanisms contributing to thrombocytopenia during DENV infection.

## Introduction

Dengue is a mosquito-borne viral disease caused by the dengue virus (DENV), a member of the *Flaviviridae* family, representing a major global public health challenge. DENV is a single-stranded, positive-sense RNA (+ ssRNA) virus, composed of a 11 kb genome that encodes a proteome comprising three structural (envelope, capsid, membrane) and seven non-structural proteins (NS1, NS2A, NS2B, NS3, NS4A, NS4B, NS5)(1). According to the World Health Organization (WHO), an estimated 390 million individuals are infected annually, and approximately 96 million develop clinical manifestations, with severe cases carrying a markedly higher risk of mortality in the absence of timely intervention(2). In India, dengue remains endemic, with 2,33,519 of reported cases and a 0.13% case fatality rate in recent surveillance data by the National Center for Vector-Borne Disease Control (NCVBDC)(3). Clinically, dengue ranges from mild febrile illness to severe, potentially fatal manifestations like dengue hemorrhagic fever (DHF) and dengue shock syndrome (DSS), characterized by vascular leakage, hemorrhage, and multi-organ dysfunction (4). Despite its substantial global burden, no specific antiviral therapy exists, with management primarily focused on supportive care, underscoring the need for novel therapeutic targets.

One of the hallmark features of dengue, closely linked to disease severity, is “thrombocytopenia”- a hematological condition defined as a platelet count below 150,000/μL, with severe cases often dropping below 50,000/μL. This reduction in platelet count results from two principal mechanisms: first, impaired platelet biogenesis due to dysregulated bone marrow megakaryopoiesis and, second, increased peripheral platelet destruction or clearance. The former is determined by significant defects in megakaryopoiesis (5). Megakaryopoiesis, the process by which platelets are generated from hematopoietic stem cells (HSCs), is regulated by a network of megakaryopoietic genes essential for normal megakaryocyte development and platelet maturation. These genes govern the different stages of platelet biogenesis: GATA1, RUNX1, and FOG-1 drive early lineage commitment of HSCs; FLI1, TAL1, and ETV6 promote megakaryoblast proliferation and maturation; while NFE2, MYH9, MYL9, MYH10, ITGA2B, and ITGB3 mediate proplatelet formation and platelet release (6). Disruption of these genetic programs, whether by altered expression or pathogenic variants, can impair successive stages of platelet formation, contributing to thrombocytopenia. Indeed, at least 40 megakaryopoietic genes and their mutations have been identified to cause diverse forms of inherited thrombocytopenia, emphasizing their indispensable role in maintaining platelet homeostasis (7).

A multitude of pathological conditions manifest thrombocytopenia as a key characteristic. Inherited forms often arise from transcription factors essential for megakaryocyte maturation. For instance, genetic variants of RUNX1 lead to familial platelet disorder, accompanied by an increased risk of acute myeloid leukemia, and are characterized by impaired megakaryocyte differentiation and reduced platelet counts (8). Likewise, mutations in GATA1 give rise to X-linked thrombocytopenia and congenital erythropoietic porphyria,(9) while alterations in c-MPL lead to amegakaryoblastic thrombocytopenia marked by complete absence of megakaryocytes(10). Deleterious mutations in the MYH9 gene, encoding cytoskeletal proteins, result in an autosomal dominant disorder presenting with platelet macrocytosis and functional defects, causing congenital thrombocytopenia (11). In addition, immune-mediated mechanisms contribute significantly, as glycoproteins and their complexes such as GPIIb/GPIIIa and GPIX/V are frequent targets of autoantibodies in disease conditions like immune thrombocytopenic purpura, leading to accelerated platelet destruction and clearance (12). Collectively, these multiple molecular pathways underscore the multifaceted mechanisms driving thrombocytopenia, with megakaryopoietic genes and immune regulation at the core. Based on these common characteristics, it is likely that dengue virus-induced thrombocytopenia is mediated through perturbations of related megakaryopoietic genes and transcription factors, along with immune-related mechanisms, highlighting the importance of investigating these molecular targets for potential therapeutic development.

Within the transcriptional network that governs megakaryopoiesis, RUNX1 (Runt-related transcription factor 1) acts as a pivotal central regulator. A study by Okuda et al. demonstrated that RUNX1 deficiency results in embryonic lethality during mid-gestation, underscoring its indispensable role in definitive hematopoiesis(13). Structurally, RUNX1 contains a highly conserved DNA-binding domain, the runt homology domain (RHD), located near the N-terminus and spanning approximately 128 amino acids. This domain enables sequence-specific DNA binding and mediates interaction with the core-binding factor subunit β (CBFβ), which stabilizes the transcriptional complex(14). In contrast, the C-terminal domain, though less conserved, is critical for recruiting transcriptional co-regulators and comprises of an activation domain, an inhibitory domain, a proline-rich region, and a nuclear localization signal (15). Functionally, RUNX1 extensively controls the transcriptional program of megakaryopoiesis by regulating genes involved in megakaryocyte polyploidization, proplatelet formation, granule biogenesis, and platelet activation. It activates the expression of platelet-specific genes like platelet factor 4 (PF4), protein kinase C theta (PRKCQ), nuclear receptor subfamily 4 group A member 3 (NR4A3), and myosin light chain 9 (myl9) (16). Conversely, it represses genes such as non-muscle myosin heavy chain IIB (MYH10), whose downregulation is essential for the transition into the endomitotic stage. Nevertheless, the compensatory crosstalk between MYH9 and MYH10 during these stages remains largely uncharted, representing an underexplored dimension of cytoskeletal regulation. Moreover, RUNX1 regulates MYH9 and members of the ETS transcription factor family, thereby coordinating cytoskeletal remodeling and platelet signaling cascades (17) (18) (19). Through these regulatory networks, *RUNX1* also drives α-granule biogenesis and cytoskeletal reorganization, both of which are crucial for platelet aggregation and spreading. Loss-of-function mutations in RUNX1 impair megakaryocyte maturation, reduce polyploidization, disrupt granule formation, and compromise platelet signaling. Clinically, these heterozygous mutations give rise to familial platelet disorder with a predisposition to myeloid malignancies (FPDMM), characterized by thrombocytopenia, bleeding diathesis, and elevated risk of leukemia (20). Given its central role in megakaryopoiesis, perturbations in RUNX1-mediated transcriptional control may substantially contribute to thrombocytopenia observed across diverse pathological conditions, including viral infections such as dengue, where platelet biosynthesis is profoundly impaired. A comprehensive understanding of the structural and functional roles of RUNX1 is therefore critical for elucidating the molecular mechanisms behind thrombocytopenia.

As an essential element in the cytoskeletal machinery of megakaryocytes, proplatelets, and platelets, MYH9 is a key determinant of platelet morphology and biosynthesis by facilitating cytoskeletal reorganization and proplatelet formation. The MYH9 gene, located on chromosome 22q12.3, spans approximately 10.6 kb, comprises 41 exons, and encodes non-muscle myosin heavy chain IIA (NMHC-IIA), a 1,960-amino-acid protein expressed ubiquitously across eukaryotic cells (21,22). NMHC-IIA forms a hexameric complex composed of two heavy chains (HCs), two regulatory light chains (RLCs), and two essential light chains (ELCs), known to function by binding to F-actin and generating contractile force through ATP hydrolysis. Structurally, the MYH9 protein consists of an N-terminal motor (head) domain with F-actin and ATP-binding/ATPase sites, a neck domain that associates with light chains, and a C-terminal tail domain containing a coiled-coil region that mediates dimerization and filament assembly as well as elements that regulate myosin activity (23,24). Based on the structural framework of MYH9, it modulates essential processes. Functionally, MYH9 supports organelle organization, membrane dynamics, and cytoskeletal remodeling in megakaryocytes, thereby promoting efficient platelet production (25). Additionally, it maintains megakaryocyte morphology via cytoskeletal contractility and extracellular matrix anchorage, facilitates the development of the demarcation membrane system (DMS) and peripheral zone, and drives proplatelet bending and branching, which are indispensable for platelet release. Consistent with these roles, MYH9 deficiency or pathogenic variants lead to reduced DMS, absence of the peripheral zone, abnormal organelle distribution, disrupted F-actin architecture, ultimately resulting in morphologically abnormal platelets and macrothrombocytopenia with enlarged, dysfunctional platelets (26). MYH9 activity is further controlled by the RHOA-ROCK signaling cascade, illustrating its indispensability for proper platelet biosynthesis(27). Collectively, these findings establish MYH9 as a crucial determinant for maintaining megakaryocyte architecture, ensuring stable anchorage to the extracellular matrix, regulating DMS formation, and maintaining the integrity of peripheral zones. Through these roles, MYH9 provides the membrane framework and cytoskeletal rearrangements necessary for normal proplatelet formation and the generation of morphologically intact, functionally competent platelets. Importantly, the MYH9 protein is known to sustain viral infection by acting as a cell-surface receptor that facilitates viral entry into host cells (28). Interestingly, this MYH9 protein has a dual regulatory mechanism to modulate viral infection. The recent study on IAV infection reveals the biopotential nature of this protein to control influenza infection, where it acts as both an anti-viral and a pro-viral factor depending on the stage of viral infection (29). However, the specific molecular checkpoints by which MYH9 governs these transitions remain insufficiently characterized, highlighting a critical gap in our mechanistic understanding.

The mechanistic basis of thrombocytopenia linked to DENV-induced impairment of megakaryopoiesis remains incompletely defined. Given the essential role of MYH9 in platelet production [24–26] and evidence that RUNX1 coordinates cytoskeletal/myosin gene programs during megakaryopoiesis (18), we interrogated the MYH9 promoter in silico and identified three consensus RUNX1-binding motifs. We therefore hypothesize that MYH9 is a direct transcriptional target of RUNX1 and that dengue virus perturbs this RUNX1-MYH9 regulatory axis, resulting in defective megakaryocyte differentiation and maturation, ultimately leading to thrombocytopenia observed in severe dengue.

## Methods and Materials

### Cell lines and virus propagation

Human hepatoma cells (*Huh7*) were cultured in Dulbecco’s Modified Eagle Medium (DMEM) high glucose media (Lonza, Himedia), containing 10% FBS (Gibco) (v/v) and 1% penicillin/streptomycin antibiotic, at 37°C in an atmosphere with 5% CO2 and 95% oxygen.

K562 cells, a myelogenous leukemia cell line, were obtained from the American Type Culture Collection (ATCC) and cultured at 37◦C, 5% CO2 in Iscove’s modified Dulbecco’s medium (IMDM) supplemented with Penicillin (100 U/ml), Streptomycin (0.1 mg/ml) and 10% Fetal Bovine Serum (FBS).

The DENV serotype 2 strain P23085 INDI-60 (Accession no. KJ918750) was propagated in the C6/36 cell line in L15 media (Himedia) supplemented with 10% FBS (v/v) and 1% penicillin/streptomycin and incubated at 28°C without CO2. Cultured supernatant was collected by adjusting the FBS concentration to 2% of the total volume. DENV supernatant was then further stored at −80°C until use.

### ChIP Assay

1 × 10^6^ *Huh7* cells were seeded in 2 separate 60 mm discs. One disc was infected with DENV-2, while another served as a control. Both discs were then incubated for 24 hours; following this, the cells were harvested, pelleted, and resuspended in 1ml ChIP Lysis buffer and incubated on ice to facilitate cell lysis. Cell lysate was centrifuged (10,000 × g for 2 min at 4°C) to isolate nuclei. Isolated nuclei were crosslinked with 1% formaldehyde to fix protein-DNA complexes. To shear chromatin, nuclei were resuspended in 1% SDS Lysis buffer and subjected to sonication (10 pulses of 45 sec on and 30 sec off) using a Bio-Rupture pulse sonicator device.

To facilitate immunoprecipitation, A/G Magnetic beads (Pierce, Cat no. 88802) were incubated with Anti-RUNX1 antibody for 2hrs. The beads were precleared by three stringent washes with PBST, then the lysate was added to the antibody-bound beads and incubated. The used lysate was removed after incubation, followed by three washes with PBST. The immunoprecipitated complexes were resuspended in ChIP Lysis buffer, and chromatin was purified with the phenol-ethanol DNA isolation method. The extracted DNA was then amplified to detect the MYH9 promoter region via RT-PCR. The immunoprecipitated RUNX1 from protein-DNA was detected using western blotting.

A similar protocol was followed for PMA-induced K562 cells. 1x10^6^ cells were seeded in 2 separate 60 mm discs. For control samples, cells were treated with 50nM PMA to induce megakaryocytic differentiation. In contrast, the DENV-2-infected samples were exposed to DENV followed by PMA induction. The K562 cells were incubated under standard conditions to allow post-infection effects to manifest. Subsequent steps for control and DENV-infected samples were the same as those described for Huh7 cells, ensuring consistency in experimental conditions.

### Dual Luciferase Reporter Assay to assess MYH9 promoter activity

The wild-type MYH9 promoter was cloned into the pGL3 Basic (Promega) Luciferase reporter vector. The MYH9 promoter construct and Renilla firefly construct were co-transfected into pre-seeded Huh7 and PMA-induced K562 cells (0.2×10^6 cells) in a 6-well plate, each separately. After 24 hours post-transfection, the cells were harvested in 1X Passive Lysis Buffer. The Assay was carried out using the Dual Luciferase Reporter System (Promega cat no. E1920).

### RNA Isolation and cDNA preparation

RNA was extracted from cultured *Huh7* and K562 cell samples using the TRIzol method. Following this, chloroform was added to the cells collected in TRIzol reagent, then after proper mixing and centrifugation, the clear aqueous layer was collected in vials, and RNA was isolated, followed by cDNA preparation using a standard protocol as mentioned previously(18)

### RT-PCR

PCR was used to analyze gene expression in Huh7 and K562 cells cultured under different conditions. Experiments were performed using cDNA prepared from the samples of both cell lines. PCR reactions were prepared on ice using Phire DNA polymerase (Thermo) containing the following components: 5x reaction buffer, 10 mM dNTPs, 20 μM forward and reverse primers, Phire DNA polymerase (5 U/μl), and cDNA template. The cycling parameters consists of initial denaturation at 98 for 3 minutes followed by amplification for (25-35 cycles) with denaturation at 98 for 30 seconds, annealing at 50-58°C for 20 seconds, extension at 72°C for 30 seconds. Lastly, a 5-minute extension at 72 °C. The reaction was then held at 4°C until further processing. Beta-Tubulin and Beta-Actin were used as endogenous controls for normalization purposes. The band intensity of samples was normalized with β-Tubulin or Beta-Actin, and graphs were plotted to check the expression level.

### Quantitative real-time-polymerase chain reaction (qRT-PCR)

qRT-PCR experiments were performed to analyze gene expression for Huh7 and K562 cultured cells under different conditions by using SYBR Green master mix (Applied Biosystems) according to the manufacturer’s protocol. For each sample, a reaction cocktail was prepared on ice containing 2x SYBR Green mix, forward and reverse primers, M-MLV reverse transcriptase (Promega), and 50 ng of RNA template from cell lysates, which was used as input for each reaction. RNA was extracted from samples with different test conditions. The cycling parameters were as follows: with first step involving cDNA synthesis at 42°C for 30 minutes, initial denaturation with hold of 95°C for 3minutes followed by a total of 40 amplification cycles comprising denaturation at 95°C for 30 seconds, annealing at 55°C for 15 seconds, extension at 72°C for 30 seconds. A melting curve analysis was done at the end of the reaction to verify amplification specificity. Beta-Tubulin and Beta-Actin serve as endogenous controls for normalization. Lastly, relative gene expressions were calculated using the 2^(-ΔΔCt) method. All qRT-PCR analyses were repeated in triplicate.

### siRNA mediated knockdown of RUNX-1 and MYH9 (*Huh7* cells and k562)

For siRNA-mediated knockdown in Huh7 cells, cells were seeded in a 12-well plate. After preparing the transfection mixture with serum-free Opti-MEM, siRNA targeting RUNX1(Santa Cruz cat no. sc-37677) and MYH9 (Santa Cruz cat no.sc-61120), and Lipofectamine RNAi-max (Invitrogen). The mixture was gently mixed and incubated at room temperature for 30 minutes to allow the formation of the siRNA-Lipofectamine complex. Following the incubation period, both the transfection complex and the Huh7 cells were carefully added to the wells. The plate was then incubated at 37°C with 5% CO□. Following the incubation period, cells for RNA analysis were harvested in TRIzol for total RNA extraction at various time points, and cells for protein analysis were harvested in RIPA buffer for western blotting. The siRNAs used in this study were sourced from Synbio Technologies.

For siRNA-mediated knockdown in K562 cells, cells were seeded in a 12-well plate. The transfection mixture was prepared with serum-free Opti-MEM, siRNA targeting RUNX1 and MYH9, and Lipofectamine RNAi-max (Invitrogen). The transfection complex was carefully added to the wells containing the seeded K562 cells. Following this, K562 cells were induced with 50 nM phorbol 12-myristate 13-acetate (PMA) to facilitate megakaryocytic differentiation. After three days of PMA induction, cells for RNA analysis were harvested in TRIzol for total RNA extraction at various time points, and cells for protein analysis were harvested in RIPA buffer for western blotting.

### Immunofluorescence Assay for analyzing the cytoskeletal changes upon DENV infection

For immunofluorescence analysis of MYH9 expression, the experiment was performed in two cell lines, Huh7 and K562. The cells were treated with DENV-2 at 5 MOI, while the control group remained uninfected. After 24 hours post-infection, cells were fixed with 4% PFA for 15 minutes, then blocked with 2% BSA in the dark at room temperature to prevent nonspecific antibody binding, and further processed for indirect immunofluorescence assay. Primary antibody incubation was then performed using an anti-MYH9 antibody to detect endogenous MYH9 expression. This was followed by incubation in the dark with an Alexa Fluor-conjugated anti-rabbit secondary antibody to prevent photobleaching. Finally, immunofluorescence images were captured using a confocal microscope to assess MYH9 localization and expression in DENV-infected and control samples.

### Western Blot Analysis

In Huh7 cells and K562 cells, lysates were collected at the indicated time points and lysed in RIPA buffer. Samples were processed with a standard protocol (18). The blot was then developed using an ECL reagent (Thermo). Primary antibodies employed in this study included Anti-MYH9 (Invitrogen), Anti-RUNX-1 (Santa Cruz), Anti-NS1 (Invitrogen), Anti-GAPDH (Abbkine) and Anti-β-tubulin (Abbkine). Secondary antibodies employed are anti-Mouse (Bio-rad: 170-6516) and anti-Rabbit (Pierce: 31460).

### Optimization of the PMA-induced K562 differentiation model

K562 cells, when induced to differentiate into megakaryocytes using phorbol esters such as PMA (Phorbol 12-myristate 13-acetate), undergo various cytological and morphological changes. Upon Phorbol-12 Myristate-13 Acetate (PMA) supplementation, cells stopped proliferating, increased in cell size and nuclear-to-cytoplasm ratio, increased attachment to the substratum, enlarged in size, and underwent endomitosis to form polyploid nuclei.

To facilitate megakaryocytic differentiation in K562 cells, cells were treated with 50nM Phorbol 12-myristate 13-acetate (PMA). The cells were incubated at 37 °C in 5% CO2 culture conditions for 0, 3, 6, and 9 days to observe differentiation characteristics under the microscope.

### Cloning of MYH9 Promoter into pGL3-Basic-Vector

Cloning was performed by amplifying the target DNA using a polymerase chain reaction (PCR). PCR products were analyzed by agarose gel electrophoresis and purified using a gel extraction kit. Following this, the purified PCR product and vector were digested with restriction enzymes (HindIII and KpnI) in a 20 µL reaction. After this, the digested DNA fragments were purified and ligated using T4 DNA ligase in a 10 µL reaction, maintaining an insert-to-vector molar ratio of 3:1 or 5:1. The ligation mix was incubated overnight at 16°C. Lastly, the ligated products were transformed into competent *E. coli* cells (DH5-Α) by heat-shocking at 42 °C for 90 seconds, immediately transferred onto ice for 5 minutes, and then recovered in LB medium. Transformants were plated and selected through PCR and restriction digestion screening. Details of the cloning are provided in supplementary figure S1.

### RUNX-1-WT and MYH9-WT plasmids procurement

pRK5-mEGFP-RUNX1-WT was a gift from Denes Hnisz (Addgene plasmid # 194574; http://n2t.net/addgene:194574; RRID:Addgene_194574) (30).

### Propidium Iodide Staining for Polyploidy Analysis using Flow-Cytometry

K562 cells were seeded in a 6-well plate and induced with 50nM PMA. Similarly, cells were seeded and treated with DMSO vehicle control samples to compare K562 cell differentiation towards the megakaryocyte lineage. The cells were incubated at 37°C in 5% CO2 culture conditions at 0, 3, and 6 days. Further, cells were harvested by centrifuging. Followed by two rounds of stringent washing of the obtained cell pellet using ice-cold 1xPBS. After washing, cells were fixed with 70% ethanol and incubated at 4°C for 30 minutes. Following that, cells were washed with ice-cold 1x PBS and treated with 200 µg/ml RNase A at 37°C for 30 minutes. After washing, cells were stained with propidium iodide at a concentration of 50µg/ml. Stained cells were resuspended in 1× PBS for flow cytometric analysis.

### Giemsa staining

For Giemsa Staining, confluent cultures of K562 cells were seeded in 60-mm Petri dishes and supplemented with 50 nM PMA. For DENV-infected PMA-induced cells, they were first infected with DENV-2, followed by 50 nM PMA induction. The cells were incubated at 37 °C in 5% CO2 culture conditions for 0, 3, 6, and 9 days. For staining the 0-day cells, cells were centrifuged at 1000 x RPM for 3 mins. The pellet was obtained and fixed in 100% methanol. The fixed cells were stained with Wright-Giemsa Staining Solution for 15-20 minutes, then washed with 1 x PBS. Cells were air-dried and mounted on a slide with mounting medium for observation under the optical microscope. Similarly, cells were harvested at 3, 6, and 9 days for staining and microscopic observation.

### Transmission Electron Microscopy

K562 cells were maintained at 37◦C, 5% CO2 in Iscove’s modified Dulbecco’s medium (IMDM) supplemented with Penicillin (100 U/ml), Streptomycin (0.1 mg/ml), and 10% Fetal Bovine Serum (FBS). The following experimental groups were taken for transmission electron microscopy analysis: (a) PMA-treated K562 cells (Mock cells): for this, 1 × 10^6 cells were seeded in a 60 mm disc. Then the cells were treated with 50nM PMA to induce megakaryocytic differentiation. (b) DENV PMA Sample: For the DENV-2-infected samples, 1 × 10^6^ cells were seeded in a 60 mm disc and exposed to DENV for 4 hours, with the virus removed before PMA induction. (c) siR-PMA (siRUNX1 KD+PMA): In this, 1 × 10^6^ cells were seeded in 60 mm dishes, then a transfection mixture with serum-free Opti-MEM, siRNA targeting RUNX1 (siRUNX1), and Lipofectamine RNAi-max (Invitrogen) was prepared and incubated at RT for 30 minutes to allow the formation of the siRNA-Lipofectamine complex. Following the incubation period, the transfection complex was carefully added to the disc containing the seeded K562 cells. Following this, K562 cells were treated with 50 nM phorbol 12-myristate 13-acetate (PMA) to induce megakaryocytic differentiation and incubated for 3 days. Similarly, the (d) siM-PMA (siMYH9 KD+PMA) samples were prepared using siRNA targeting MYH9 (siMYH9), and a similar protocol was used. Then, the remaining rescue samples (e) siRUNX1+ p-GFP-RUNX1 and (f) siMYH9+ p-V5-MYH9: In this, 1 × 10^6^ K562 cells were seeded in 60 mm dishes and transfected with siMYH9 or siRUNX1 at a 20nM concentration. Twelve hours post-transfection, the respective KD cells were transfected with p-GFP-RUNX1 or p-V5-MYH9 plasmid constructs at a concentration of 1 μg. Then, 7-8 hours post-transfection, the cells were treated with 50 μM PMA and incubated for 3 days before proceeding with the TEM protocol. The TEM protocol was as follows: cells from the above samples were trypsinized and washed with 0.1 M PBS to remove cell culture medium. Followed by primary fixation with (1.5% glutaraldehyde, 1.25% paraformaldehyde, and 0.1 M phosphate buffer, pH 7.0) for 15 minutes at RT and resuspended in 0.1 M PBS following this post fixation was done using 1% osmium tetroxide (OsO_4_) for 1 hour at RT and then were washed twice using distilled water. After this, 1% uranyl acetate was added to each sample, and the samples were incubated for 1 hr at RT. Next, the samples were dehydrated by incubating using a series of gradually increasing concentrations of ethanol (30%, 50%, 70%, 80%, 90%, 95% and 100%) for 10 minutes each, followed by two additional incubations of 10 minutes each in 100% ethanol from a freshly opened bottle. The samples were then embedded in resin by infiltrating them with 3:1, 1:1, and 1:3 embedding resins for 2 h at each ratio. Subsequently, the samples were incubated overnight with 100% fresh resin. Following this, the cells were transferred into BEEM capsules (EMS) and polymerized at 70°C for 4 days. Then ultrathin sectioning was done with a diamond knife (Diatome) on an ultramicrotome (Ultracut UCT; Leica Microsystems), collected on 200 mesh copper grids (EMS), poststained with uranyl acetate and lead citrate, and visualized with a Talos S200 transmission electron microscope (TEM; Thermo Fisher Scientific), operating at 200kV.

### Statistical Analysis

The data were represented as mean ± SD (standard deviation) derived from independent experiments. Significant Differences between the two groups were assessed using an unpaired two-tailed t-test, and differences among more than two groups were assessed using analysis of variance (ANOVA) in GraphPad Prism (GraphPad Software Inc., San Diego, CA). Statistical significance was defined as p<0.05.

## Results

### Gene expression profiling of a subset of Megakaryopoitic genes in dengue-infected *Huh7* cells

Thrombocytopenia, low platelet count, is a key clinical feature of Dengue virus pathogenesis. To investigate the cause of decreased platelet count during dengue infection, we sought to decipher the expression of genes that participate at various stages of platelets biogenesis/megakaryopoisis. To achieve this, *Huh7* cells were infected at MOIs of 1, 5, and 10, and samples were harvested 24 hours post-infection. The expression of megakaryopoitic genes was assessed using RT-PCR. Mock-treated cells were taken as a control. A subset of megakaryopoitic genes were analyzed, such as CYCS, DIPH1, MASTL, SRC, ETV6, ACTN1, ANKRD26, TRPM7, ABCG5, RUNX-1, RBM8A, and MYH9 in *Huh7* cells infected with DENV (primer sequences have been provided in supplementary Table 1). Our data suggest a significant, gradual reduction in MYH9 expression during DENV infection (Figure 1A). MYH9, a non-muscle myosin II A protein, is a cytoskeletal protein that is required for cell motility and cell shape. Importantly, platelets exclusively produce the MYH9 isoform. MYH9 protein is required at the maturation stage of platelet biogenesis, particularly during platelet release from proplatelets.

**Figure 1.**
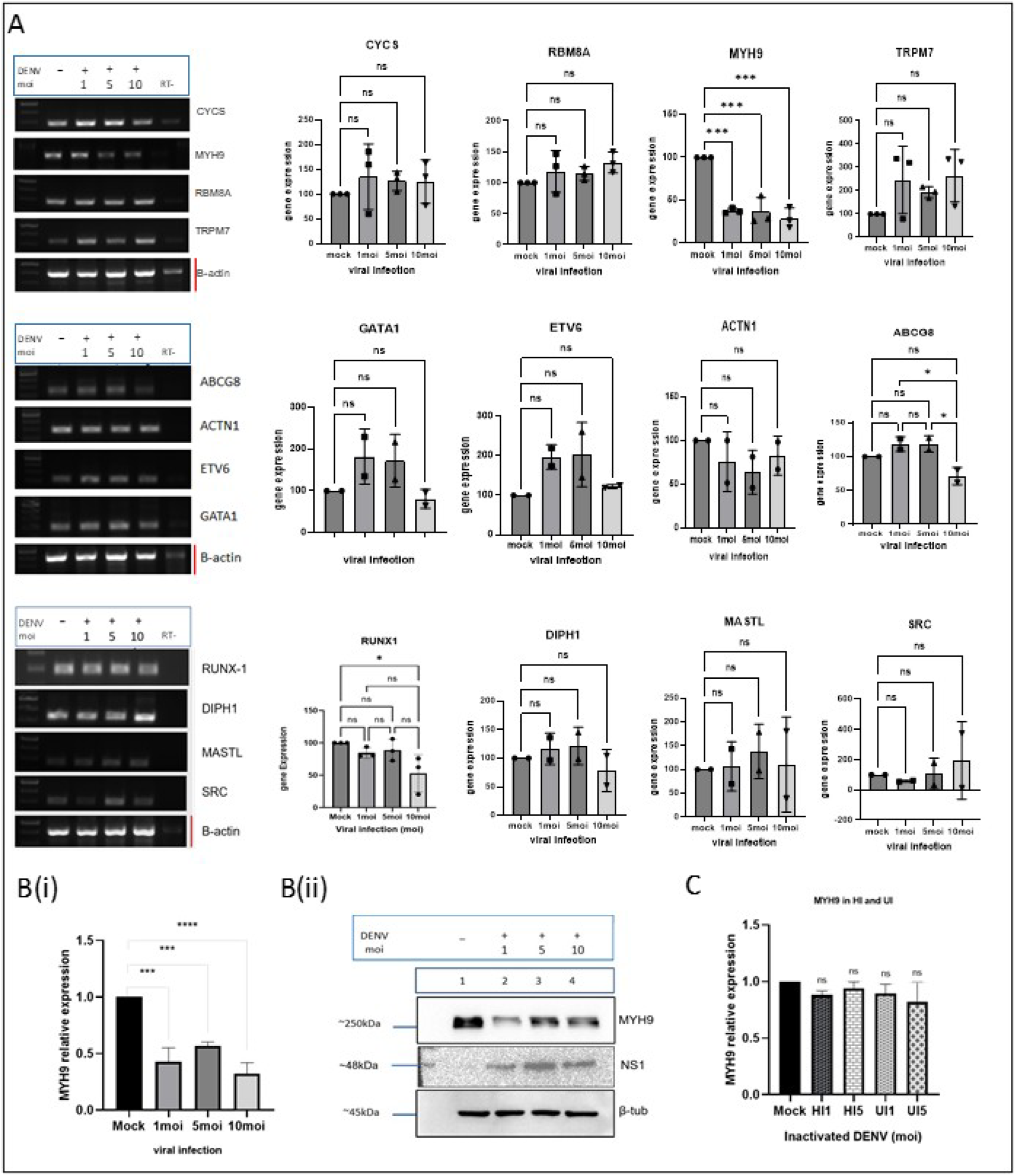
Gene expression profiling of a subset of Megakaryopoitic genes in dengue-infected *Huh7* cells. *Huh7* cells were infected with Dengue Virus at MOIs of 1, 5, and 10, followed by cell lysate preparation and differential expression analysis of Megakaryopoetic genes. (A) RT-PCR analysis shows differential expression of genes involved in platelet biogenesis during DENV infection. (B)(i) qRT-PCR and (ii) Immunoblotting analysis show the significant down-regulation of the MYH9 gene during DENV infection. Along with detection of dengue virus NS1 protein. (C) RT-PCR analysis of the MYH9 gene expression from heat-inactivated and UV-inactivated dengue-infected samples shows non-significant changes.

Additionally, we further investigated the MYH9 gene expression at the RNA level and protein level using qRT-PCR and the western blotting technique, respectively (Figure 1B (i) and (ii)). The data indicate a decrease in MYH9 gene expression at both the RNA and protein levels. The viral detection from qRT-PCR data is shown in Supplementary Figure S2. To further assess the specificity of this observation, we analyzed MYH9 gene expression in heat- or UV-inactivated dengue-infected Huh7 cells. The RT-PCR results indicate no significant change in MYH9 expression during inactivated dengue virus infection (Figure 1C). As is known, cytoskeletal remodeling is fundamental for platelet production; this progressive downregulation of MYH9 gene expression may contribute to impairment of megakaryopoiesis, thereby aligning with “Thrombocytopenia,” a hallmark feature of dengue. Taken together, we hypothesize that MYH9 is an important protein required for platelet maturation and decreased expression of the MYH9 gene may lead to reduced platelet synthesis.

### RUNX-1 mediated transcriptional regulation of MYH9 is perturbed during DENV infection in *Huh7* cells

Next, we examine the regulation of MYH9 gene expression under DENV pathogenesis. Consequently, we performed *in-silico* analysis using the Eukaryotic Promoter Database to list the transcription factor binding sites in the promoter region of the MYH9 gene. We selected the RUNX-1 transcription factor, which has three potential binding sites in the MYH9 promoter region (Figure 2A). To substantiate RUNX-1-mediated MYH9 gene expression regulation, we investigated whether RUNX-1 interacts with the MYH9 promoter. To achieve this, we implemented a ChIP-based workflow to assess RUNX-1 binding in uninfected mock cells and DENV-infected cells. *Huh7* cells were seeded in 60-mm dishes at a density of 100 × 10^4^ cells. Twenty-four hours post-infection, the cells were fixed in 1% formaldehyde followed by sonication and DNA extraction. The RT-PCR analysis of ChIP DNA of mock-treated samples revealed a strong interaction between RUNX-1 TF and the MYH9 promoter region in mock-treated cells, whereas a significant reduction ∼80% was observed upon DENV infection (Figure 2B).

**Figure 2.**
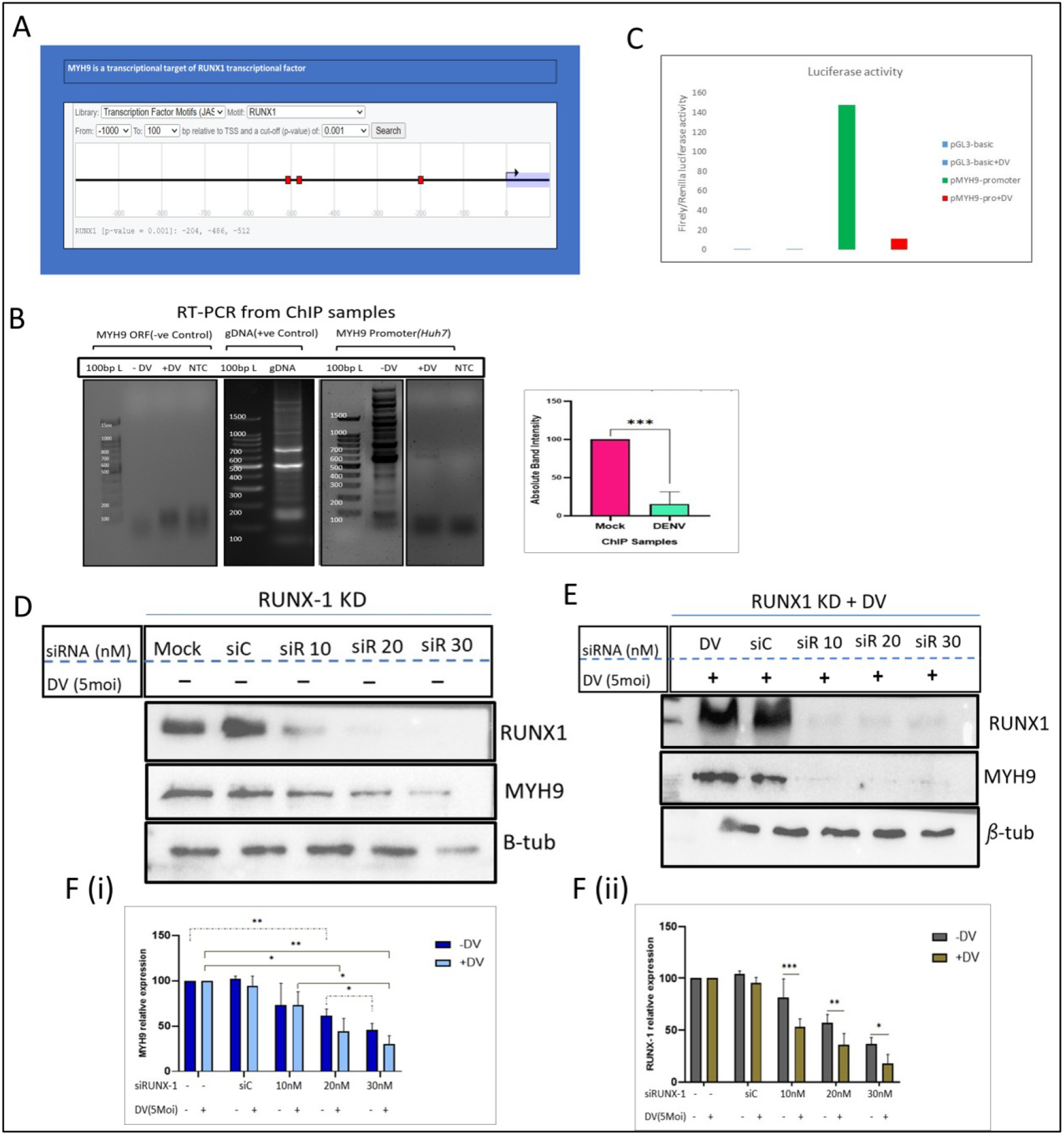
RUNX-1-mediated transcriptional regulation of MYH9 is perturbed during DENV infection in *Huh7* cells. RUNX-1-mediated MYH9 gene regulatory axis has been evaluated in the DENV-infected *Huh7* cell line. (A) *In-silico* analysis from the Eukaryotic Promoter Database (EPD) indicates the three potential binding sites for the RUNX-1 transcription factor in the MYH9 gene promoter region. (B) MYH9-promoter DNA was precipitated with RUNX-1 antibody in vitro, and the precipitated DNA was subjected to RT-PCR analysis. ChIP data suggest a decrease in the physiological association between the MYH9 promoter and the RUNX-1 transcription factor during dengue virus infection. (C) *Huh7* cells were transfected with an empty pGL-3-basic vector as a vector control (VC) at a concentration of 1 µg or with a plasmid carrying MYH9-promoter (pMYH9-promoter) at a concentration of 1 µg. Twenty-four hours post-transfection, cells were infected with DENV at a 5 MOI, and cells were harvested 24 hours post-infection in Passive Lysis Buffer (PLB). The cells were subjected to a luciferase assay. The analysis indicates a decrease in MYH9 Promoter activity during DENV infection. (D) Huh7 cells were transfected with non-specific siCtrl (siC) and RUNX-1-specific siRNA at concentrations of 10nM, 20nM, and 30nM. Cells were harvested 24 hpt and were subjected to MYH9 and RUNX-1 expression analysis. Western blot results indicate that RUNX-1 knockdown cells showed a concomitant reduction in MYH9 protein levels. (E) RUNX-1 knockdown cells were infected with DENV at a 5 MOI, and samples were used to analyze the MYH9 and RUNX-1 protein abundance using immunoblotting. This shows a reduction in both proteins during dengue infection. (F)(i) and (ii) RT-PCR analysis was performed from RUNX-1 knockdown cells in the absence and presence of DENV infection to determine the expression of MYH9 and RUNX-1 compared to mock-transfected cells, respectively. RT analysis suggests that dengue infection downregulates both MYH9 and RUNX-1 gene expression.

To validate the specificity of this interaction, the MYH9 ORF was used as a negative control, as no amplification was observed in either condition, demonstrating that RUNX-1 binding is specific to the MYH9 promoter region. Additionally, a positive control was included in which genomic DNA (gDNA) successfully amplified the expected product. These investigations suggest that DENV infection perturbs the regulatory axis of the MYH9 gene by interfering with RUNX-1 binding at its promoter. Our data show that DENV infection might affect RUNX-1 occupancy on the MYH9 promoter, potentially perturbing its regulatory function.

Subsequently, after establishing that the RUNX-1 Transcription factor binds the MYH9 promoter region and that DENV perturbs this physiological interaction, we sought to determine whether this disruption affects MYH9 promoter activity. We performed a dual luciferase assay. Primarily, we cloned the MYH9 promoter region into the pGL3-Basic promoter-less vector upstream of the firefly luciferase gene. 1 µg of the pGL3-Basic-MYH9-promoter construct was co-transfected along with the pRLTK Renilla luciferase control into the *Huh7* cell line to quantitatively measure the changes in MYH9 promoter activity. After 24 hours post-transfection, the cells were infected with dengue virus at an MOI of 5. The cells transfected with the empty pGL-3-basic vector were taken as the vector control (VC). The cells were harvested 24 hours post-infection. The cells were assessed for luciferase activity. The data indicate that in p-MYH9-promoter-transfected cells, MYH9 promoter activity was significantly higher, as evidenced by strong luciferase expression. However, a significant reduction in the luciferase activity was observed upon DENV infection, suggesting that the transcriptional activity of the MYH9 promoter is repressed upon DENV infection (Figure 2C). To ensure specificity, the pGL3-Basic Vector was used as a negative control, which yielded minimal luciferase activity, thereby confirming that the observed signal was promoter-driven. Additionally, a DENV-infected pGL3-Basic vector was used as an additional control, ruling out non-specific effects of viral infection on luciferase expression.

Our findings collectively indicate that DENV infection disrupts MYH9 transcriptional regulation, likely through impaired RUNX-1 binding, which may contribute to cytoskeletal dysregulation and platelet dysfunction observed during infection.

Furthermore, to elucidate the regulatory effect of RUNX-1 on the MYH9 gene, we aim to determine how modulating RUNX-1 expression may influence MYH9 abundance. We assessed the effect of siRNA-mediated RUNX-1 knockdown on the MYH9 gene level. The expression of RUNX-1 was downregulated using RUNX-1-specific siRNA. The *Huh7* cells were transfected with non-specific siRNA (siC) and specific siRUNX-1 at final concentrations of 10 nM, 20 nM, and 30 nM. The effect of siRUNX-1 was determined 24 hours post-transfection using immunoblotting. This suggested a dose-dependent depletion of RUNX-1 protein. Additionally, MYH9 protein abundance was also assessed in RUNX-1 knockdown cells. The data suggested a concomitant decrease in MYH9 expression under RUNX-1 knockdown conditions (Figure 2D).

To study the impact of DENV infection on RUNX-1-mediated MYH9 regulation, RUNX-1-knockdown cells were infected with DENV at 5 MOI, and at 24 hpi, cells were harvested to monitor the expression of RUNX-1 and MYH9. Our western blot data show that dengue virus exacerbated the downregulation of both RUNX-1 and MYH9 (Figure 2E).

Subsequently, we performed a comparative assessment of MYH9 and RUNX-1 mRNA levels in RUNX-1 knockdown cells and DENV-infected RUNX-1 knockdown cells. In Uninfected cells, a dose-dependent downregulation of RUNX-1 mRNA expression in *Huh7* cells with increasing siRNA concentrations was observed, confirming knockdown efficacy (Figure 2F(i)). Notably, in DENV-infected samples, an additional suppressive effect on RUNX-1 expression beyond the levels observed in uninfected RUNX-1 knockdown samples was observed (Figure 2F(ii)). This suggests that DENV itself contributes to RUNX-1 suppression, either directly through viral-mediated transcriptional repression.

In parallel, we evaluated MYH9 expression; under uninfected knockdown conditions, MYH9 mRNA levels showed a proportional decrease in response to RUNX-1 knockdown. Thereby, supporting the hypothesis that RUNX-1 may directly regulate the transcription of MYH9. In DENV-infected RUNX-1-knockdown samples, the expression of MYH9 transcripts was even more decreased, suggesting that DENV perturbs the binding of RUNX-1, which thereby exacerbates the transcriptional suppression of MYH9.

These findings suggest that RUNX-1 plays a pivotal role in regulating MYH9 expression. Therefore, the depletion of RUNX-1, either through siRNA-mediated knockdown or DENV infection, results in a significant reduction in MYH9 levels. This reduction in MYH9 may contribute to cytoskeletal disruptions observed during DENV infection, highlighting a potential mechanism through which DENV alters cellular architecture and function.

### MYH9 dysregulation leads to cytoskeletal defects in the *Huh7* cell line

To investigate functional consequences of the MYH9 down-regulation on viral replication and over cell cytoskeletal integrity, we performed MYH9 knockdown in *Huh7* cells. MYH9 expression was knocked down using specific siRNA against the MYH9 gene. The *Huh7* cells were mock-transfected or transfected with non-specific siRNA (siC) and siMYH9 at a final concentration of 10nM. The cells were harvested 24 hpt and assessed for MYH9 knockdown efficiency. The western blotting results indicate efficient reduction in the MYH9 protein (Figure 3A(i)), and qRT-PCR indicates reduced RNA levels (Figure 3A(ii)). To further study the effect of MYH9 depletion on dengue virus replication, we seeded Huh7 cells and then transfected them with siMYH9. The transfected cells were then infected with DENV, followed by detection of dengue viral protein NS1 using western blotting (Figure 3B(i)). The data suggest improved detection of the viral protein in MYH9-knockdown cells. The MYH9 reduction under dengue infection has been confirmed with real-time PCR (Figure 3B(ii)).

**Figure 3.**
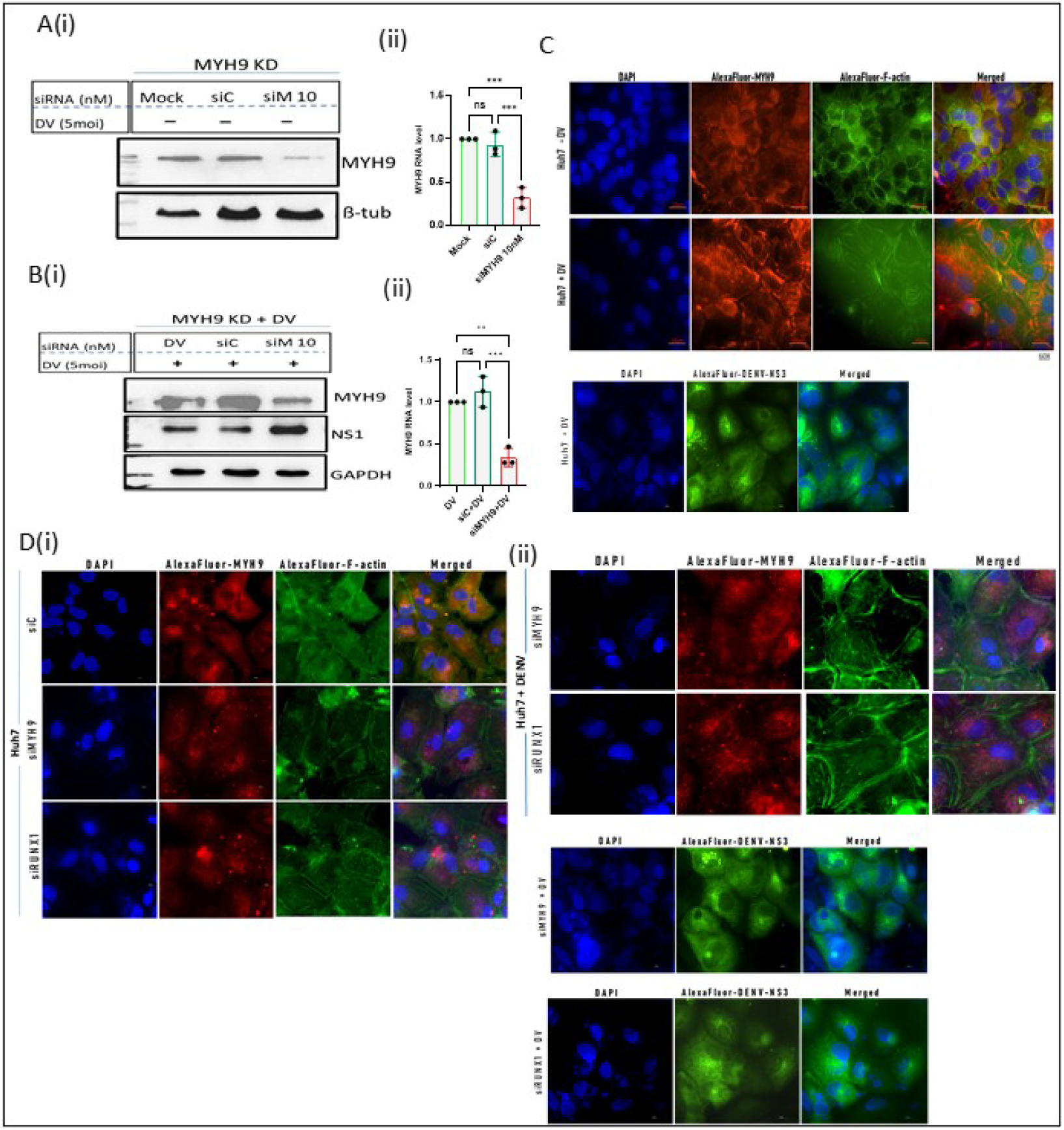
MYH9 dysregulation leads to cytoskeletal defects in the *Huh7* cell line. *Huh7* cells were transfected with MYH9-specific siRNA duplex at a final concentration of 10nM, followed by DENV infection at 5 MOI, and samples were collected at 24 hpi. (A) Cells were mock-transfected, transfected with non-specific siCtrl, and specific siRNA against MYH9 at a final concentration of 10nM. The MYH9 gene knockdown was confirmed with (i) western blotting and (ii) qRT-PCR. (B) (i)DENV infection was given to the cells 24 hpt. The cells were harvested at 24 hpi to determine the levels of Dengue virus NS1 protein relative to mock- and siC-transfected cells. The western blot analysis indicates the upregulation of viral protein NS1 in MYH9 knockdown cells, and (ii) real-time PCR analysis indicates reduced expression of the MYH9 gene under dengue infection. (C) The mock- and DENV-infected cells were subjected to an immunofluorescence assay (scale bar 10μm) using Anti-MYH9 and Alexa Fluor 488-conjugated phalloidin. The result shows that dengue virus disrupts cytoskeletal bundles, leading to disruption of the cell’s cytoskeletal structure. (D) The cytoskeletal integrity was evaluated under MYH9 and RUNX-1 knockdown in the presence and absence of dengue virus. (i) Immunofluorescence imaging from the siMYH9 and siRUNX-1 cells displays shortened, disorganized, rearranged cytoskeletal stress fibers. (ii) This effect gets more pronounced under dengue virus infection. Additionally, Dengue virus detection was confirmed with DENV-NS3 protein.

Moreover, we determined whether DENV-associated downregulation of MYH9 protein results in a functional defect in cell cytoskeletal integrity. We first aimed to compare cytoskeletal organization between mock- and DENV-infected cells. For this, an indirect immunofluorescence assay was performed in mock-treated and DENV-infected *Huh7* cells using an anti-MYH9 antibody to assess its localization and expression. *Huh7* Cells were stained with primary MYH9 antibody and secondary antibody Alexa Fluor-conjugated anti-rabbit antibody (red) to detect primary antibody anti-MYH9, and nuclei were counterstained with DAPI (blue). Alexa 488-conjugated F-actin stain was used to probe F-actin fibers (green) (Figure 3C). Upon analyzing MYH9 expression through indirect immunofluorescence Assay staining, comparable differences were observed in the cytoskeletal organization of mock-treated Huh7 cells compared to the DENV-infected *Huh7* cells. In mock-treated cells, MYH9 exhibited a well-organized, filamentous distribution, reflecting intact, structurally stable cytoskeletal bundles. The MYH9 signal appeared continuous and uniformly distributed throughout the cytoplasm, aligning with its known role in maintaining cytoskeletal integrity and cellular framework.

However, in DENV-infected cells, MYH9 localization displayed a markedly altered pattern. The staining appeared disorganized, with a fragmented distribution, suggestive of cytoskeletal disassembly. The loss of filamentous MYH9 structures, coupled with a diffuse cytoplasmic signal, indicates potential cytoskeletal destabilization and impaired structural integrity. This disruption may be attributed to viral-mediated dysregulation of MYH9, leading to compromised cytoskeletal dynamics. Such alterations could have significant implications for cellular processes, including intracellular trafficking, membrane dynamics, and overall cell morphology, further supporting the hypothesis that DENV infection perturbs MYH9-mediated cytoskeletal regulation.

Overall, in mock-treated cells, MYH9 exhibits a well-organized and uniform distribution, indicating an intact cytoskeletal structure. However, upon DENV infection, MYH9 staining appears disorganized, with a loss of filamentous structure, indicating cytoskeletal destabilization and impaired network integrity.

### Differential expression of Megakaryopoitic genes in PMA-induced K562 cells

To examine the differential expression of Megakaryopoitic genes in PMA-induced K562 cells, we re-analyzed the RNA-seq data submitted to the GEO Database (GSE186089) from Dr. Sankar Bhattacharyya’s Lab (THSTI, Faridabad, India). They have taken a total of 12 RNA samples for RNA Sequencing. The details of sample grouping are provided in Supplementary Figure S3. To interrogate expression changes in our genes of interest, we generated heatmaps showing differential expression across samples (Figure 4A). We plotted log2 fold changes for megakaryopoiesis genes in clustered heatmaps comparing all infected samples versus all uninfected samples. To create the heatmaps, we first compiled lists of all significantly differentially expressed genes associated with megakaryopoiesis. We loaded these gene lists into an R package along with the normalized expression count data for all samples. We used the heatmap or heatmaply packages to plot the expression values, setting parameters to display log2 fold changes ≥1.5 for upregulated genes, ≤0.5 for downregulated genes, and p-values ≤0.05 for each gene in each set. On each heatmap, each row represents a single gene, and each column an individual sample set. The color intensity was scaled to the fold-change magnitude to visually distinguish subtle from large expression differences. The red color indicates increased expression, while blue denotes decreased expression in infected samples compared with uninfected controls. The white color denotes non-significant gene expression in the particular set (Figure 4A).

**Figure 4.**
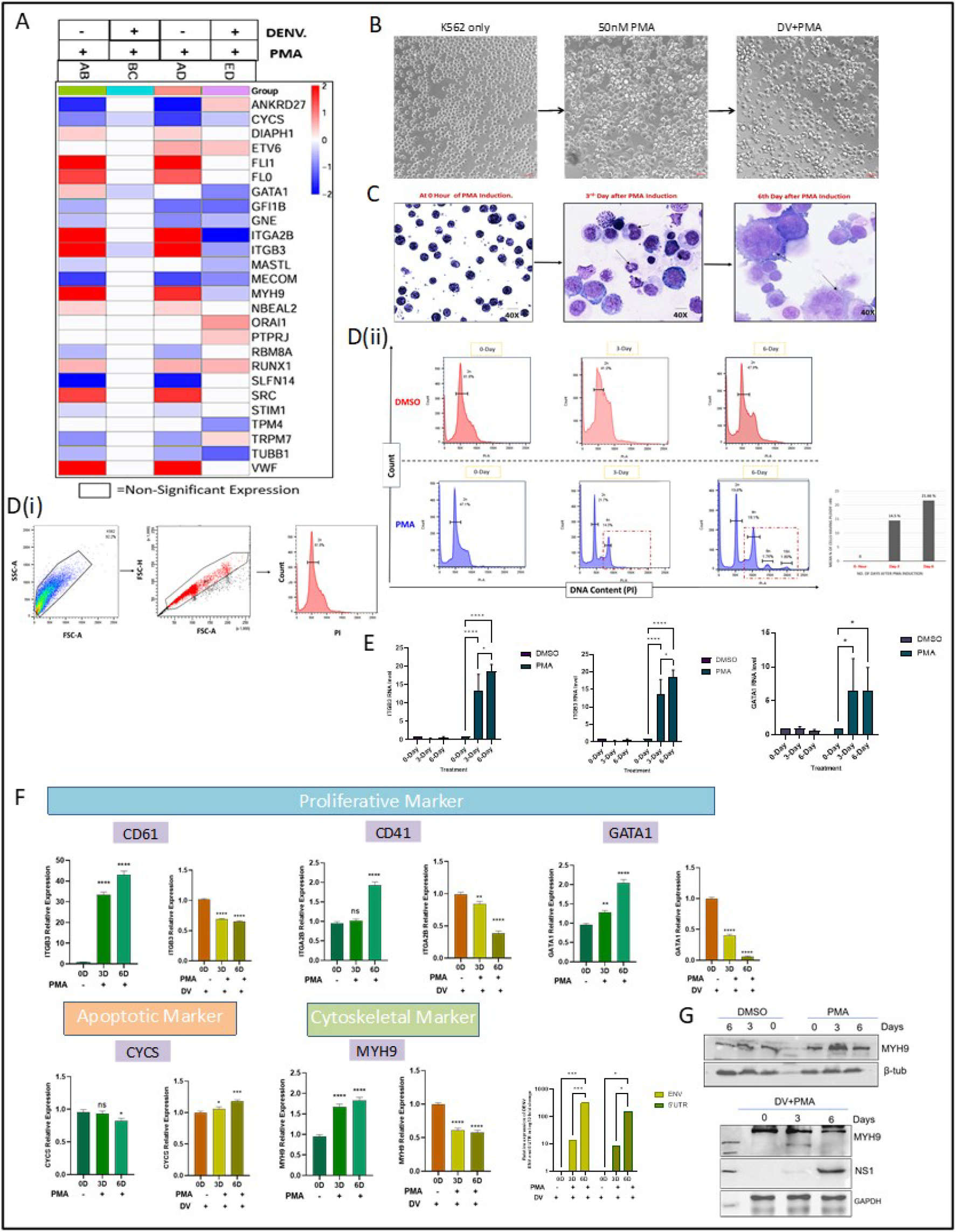
Differential expression of Megakaryopoitic genes in PMA-induced K562 cells. Megakaryopoiesis gene expression was analyzed in PMA-induced K562 Megakaryocytic cells. (A) Heatmap of differentially expressed megakaryopoitic genes, generated from RNA-seq data retrieved from the GO database GSE186089. (B) Bright-field imaging (scale bar 50μm) of K562 cells, PMA-differentiated K562 cells, and dengue virus-infected PMA-differentiated cells. The image shows the effect of PMA induction, which increases K562 cell size and adherence to the culture flask. Dengue affects PMA differentiation. (C) PMA induction was given to K562 cells at a concentration of 50 nM, and cells were analyzed for megakaryocytic differentiation at 0-, 3-, and 6-day post-induction. Giemsa staining of PMA-induced K562 cells at 0-, 3-, and 6-day post-induction shows the increase in cell size and nuclear lobulation. (D) (i) Gating strategy for flow cytometry analysis. (ii) Flow cytometry using PI staining of PMA-induced K562 megakaryocytic cells shows the increase in nuclear ploidy of the cells. (E) qRT-PCR analysis of the platelet proliferation marker genes ITGA2B (CD41), ITGB3 (CD61), and GATA from the PMA-induced K562 cells indicates the upregulation of these marker genes. The result shows megakaryocytic differentiation of K562 cells under PMA induction. (F) qRT-PCR analysis of Proliferative marker genes (ITGA2B, ITGB3, GATA), Apoptotic gene (CYCS), and cytoskeletal gene (MYH9) from PMA-K562 cells and dengue virus-infected PMA-K562 cells. The quantitative PCR results show that the dengue virus downregulates the platelet proliferative marker genes ITGA2B, ITGB3, and GATA, as well as the cytoskeletal gene MYH9. The dengue virus upregulates the apoptotic gene CYCS. (G) Immunoblotting from PMA and DV+PMA cells shows the upregulation of MYH9 in PMA samples and MYH9 downregulation in DV+PMA samples.

Since K562 cells can undergo megakaryocytic differentiation and share key transcriptional regulators with platelet development, it remains to be determined whether DENV affects these cells in a similar manner. Therefore, investigation in K562 cells could provide valuable insights into how DENV-induced transcriptional dysregulation impacts megakaryopoiesis and platelet function, ultimately leading to a hallmark symptom of DENV infection that is “Thrombocytopenia”.

K562 cells were cultured in complete IMDM media, under normal culture conditions of 5% CO2 and 37°C. These are suspension cell lines that appear round and smooth in morphology with a diameter ranging from 8-20 μm. Upon addition of phorbol 12-myristate 13-acetate (PMA) to the culture media at a concentration of 50 nM, morphological changes were observed in the K562 cells over a period of 3-6 days. After PMA exposure, many cells began to adhere and flatten out on the substratum surface. Along with an increased cytoplasmic-to-nuclear ratio, extensive pseudopodia were observed in some cells. Interestingly, dengue-infected PMA-induced K562 cells showed reduced differentiation properties compared to the K562 cells. The brightfield image shows the effect of dengue virus on megakaryocytic cell differentiation (Figure 4B).

These changes were tracked and verified by Giemsa staining, which differentially stains cellular components, allowing clear visualization of the nucleus, cytoplasm, and cell membranes under light microscopy. K562 cells were cultured in a 60 mm Petri plate, supplemented with 50nM PMA, and incubated for 2 hours (0 day), 3 days, and 6 days to observe changes due to PMA-induced differentiation. Following that, cells were then fixed and stained with freshly diluted Giemsa stain according to the protocol. Microscopic examination of K562 cells at 0 hours of PMA induction revealed a mononuclear morphology characteristic of the undifferentiated state, including large nuclei; no polyploidization, cytoplasmic maturation, or formation of proplatelets was evident. Cell size remained homogeneous with few granules observed (Figure 4C). At 3 days, cells were attached to the substratum and exhibited increased size and nuclear and cytoplasmic complexity, with lobulated or segmented nuclei visible in subpopulations, though the majority retained immature single nuclei. Moreover, some cells, as marked in Figure 4C, showed bilobed to multilobed nuclei. No clear proplatelet extensions were observed at this intermediate stage, but these characteristic features indicate that K562 cells were differentiating into megakaryocytes. At 6 days post-induction, fully differentiated megakaryocytic phenotypes predominated, characterized by large, multilobulated nuclei and expanded, mature cytoplasm with extensive granularity and vacuolations seen in Figure 4C. Cell ploidy increased, along with enhanced nuclear complexity. The morphological profile was consistent with a terminal megakaryocytic maturation stage.

By analyzing the following Giemsa staining results, we inferred that PMA-induced differentiation in K562 cells recapitulates megakaryocyte developmental features to varying degrees.

Following this, to further evaluate the polyploidization dynamics during megakaryocytic differentiation of K562 cells over 6 days of PMA induction, we performed DNA content/polyploidy analysis by propidium iodide staining and flow cytometry. Propidium iodide is a fluorescent dye that intercalates in DNA, allowing quantification of cellular DNA content. Cell populations were first distinguished from debris on a forward-scatter area (FSC-A) versus side-scatter area (SSC-A) plot, establishing an initial uniform event population for fluorescence analysis (Figure 4D (i)). Doublets were excluded by gating on singlet events using a FSC area (FSC-A) versus FSC height (FSC-H) plot. Cell cycle status and ploidy measurements were assessed using a DNA content histogram that plots propidium iodide fluorescence intensity on the x-axis. Quantification of event percentages across defined ploidy peaks (2N, 4N, 8N, 16N, 32N) was facilitated by overlaying markers on the DNA histogram, aligned to expected fluorescence intensities, in FlowJo software.

This gating strategy standardized the fluorescence quantification and facilitated consistent ploidy comparisons across treatment conditions over the time course of K562 cell megakaryopoietic differentiation. In addition to PMA induction and DMSO vehicle control, a parallel set of K562 cell cultures was treated with 20 mM sodium butyrate over the 6-day time course (supplementary figure S4). In contrast to PMA, DMSO vehicle control-treated K562 cells exhibited no appreciable emergence of polyploid sub-populations over the 6-day time course. 0 hr profiles showed the standard diploid 2N peak distribution at approximately 61.8% (Figure 4D (ii)). On day 3, minimal deviations were observed, with 2N cells persisting at 41%. By day 6, the 2N fraction accounted for 47.9%, with no significant peaks observed at 4N or higher. In undifferentiated 0 hr K562 cells, the PI histogram peak corresponded to diploid (2N). DNA content regardless of treatment, indicating a progenitor state. By day 3 of PMA induction, emergence of defined 4N peaks was detectable, comprising nearly 14.5% (Figure 4D (ii)). Further, by day 6 of PMA induction, with continued expansion of 4N, 8N, and 16N peaks, 8N and 16N peaks also appeared, comprising about 1.76% and 1.80% of cells, respectively. This signified the initiation of endomitosis and polyploidization, characteristic of megakaryocyte differentiation in K562 cells, upon PMA induction.

By plotting the mean fluorescence intensity graph (Figure 4D (ii)) for the PMA-treated K562 cell samples, we can clearly infer that the population of cells having ploidy higher than 4n increases as the days of incubation progress from 0 to 3 to 6 days.

To evaluate the gene expression profile of K562 cells undergoing PMA-induced megakaryocytic differentiation over 6 days, we assessed the expression of genes and transcription factors involved in megakaryopoiesis. The analysis included PMA-treated cells and DMSO vehicle controls at 0 hours, 3 days, and 6 days post-induction, assessed by RT-PCR. Megakaryocyte-associated genes examined included the surface receptors CD41 and CD61, as well as key lineage-determining transcription factors such as GATA1 (primer sequences are provided in Supplementary Table 1).

K562 cells were treated with phorbol 12-myristate 13-acetate (PMA) to induce megakaryocytic differentiation or with DMSO as a control. Cells were harvested at days 0, 3, and 6 of treatment. Total RNA was isolated from the cell samples. Beta-Tubulin was used as an internal control (Figure 4E).

By analyzing the expression of GATA1, ITGA2B, and ITGB3 by qRT-PCR during megakaryopoiesis in PMA-treated K562 cells, we find that these genes are significantly upregulated in response to PMA-induced megakaryocytic differentiation. This correlates with the known role of ITGA2B, ITGB3, and GATA1 as key megakaryocytic transcription factors.

Lastly, we investigated the expression profiles of several essential megakaryopoietic genes and transcription factors. The aim was to evaluate gene expression in K562 cells undergoing PMA-induced megakaryocytic differentiation over 6 days and to assess how dengue virus affects the expression of these key genes. For this experimental analysis, we included PMA-treated cells, and samples were collected at 0, 3, and 6 days post-induction for qRT-PCR analysis. The selected megakaryocyte-associated genes included the proliferative markers CD41 and CD61, the lineage-determining transcription factors GATA1 and MYH9, and the apoptotic marker CYCS. Similarly, to investigate the effect of dengue virus on the expression of these megakaryopoietic genes, we prepared another set of samples in which K562 cells were infected with dengue virus at 5 MOI, followed by PMA induction. qRT-PCR data (figure 4F) reflects a significant upregulation of ITGB3, ITGA2B, GATA1 and MYH9 transcription during PMA-induced megakaryocytic differentiation in K562 cells, as indicated by elevated mRNA levels. However, the expression of these genes is reduced in PMA-treated cells infected with DENV, indicating DENV interferes with the expression of proliferative marker genes during megakaryopoiesis. Thus, our results provide evidence that DENV modulates megakaryopoiesis by inhibiting the transcription of megakaryopoietic genes normally observed during PMA-induced K562 differentiation.

Importantly, CYCS, a critical protein involved in cellular energy production and apoptosis, also plays a role in platelet clearance; the qRT-PCR results demonstrate a marked downregulation of CYCS transcription during PMA-induced megakaryocytic differentiation, as reflected by significantly reduced CYCS RNA levels in PMA-treated K562 cells from days 3 to 6 (Figure 4F). This downregulation suggests suppression of cytochrome c-mediated apoptotic pathways, consistent with the requirement for controlled apoptosis during megakaryopoiesis.

However, in the presence of dengue virus, CYCS gene expression increases progressively over the days. Moreover, we performed immunoblotting to detect MYH9 protein levels in PMA and DENV+PMA samples. The MYH9 protein expression is increased under PMA differentiation; in contrast, dengue virus-infected PMA-differentiated cells demonstrated reduced expression of MYH9 protein (Figure 4G).

Overall, our gene expression analysis data show a multi-channel effect of Dengue infection on platelet biogenesis. DENV selectively downregulates the proliferative marker genes CD61, CD41, and GATA, the cytoskeletal gene MYH9, and upregulates the CYCS apoptotic gene to promote platelet destruction.

### RUNX-1-mediated transcriptional regulation of MYH9 is perturbed during DENV infection in the K562-megakaryocytic model

Next, we investigated the working potential of RUNX-1-mediated MYH9 gene regulation and its perturbation under DENV infection in PMA-induced K562 cells. Similarly, in K562 cells, we demonstrated the physical association between the RUNX-1 transcription factor and the MYH9 promoter region. The K562 cells were seeded, dengue infection was introduced at the time of seeding, and PMA induction was performed 2 hours post-infection. PMA-induced K562 cells without dengue virus infection were taken as a control. The cells were harvested in ChIP lysis buffer and subjected to immunoprecipitation of the RUNX1-MYH9 Promoter complex using an anti-RUNX1 antibody. DNA was extracted from this complex and used to amplify the MYH9 promoter region. The RT-PCR amplification shows greater enrichment of the MYH9 promoter region in mock cells than in dengue-infected cells. In DENV-treated samples, a significant ∼30% reduction was observed. Although the observed reduction is partial but significant, it suggests that even a slight disruption in RUNX-1 binding to the MYH9 promoter can impair cytoskeletal integrity and platelet function (Figure 5A).

**Figure 5.**
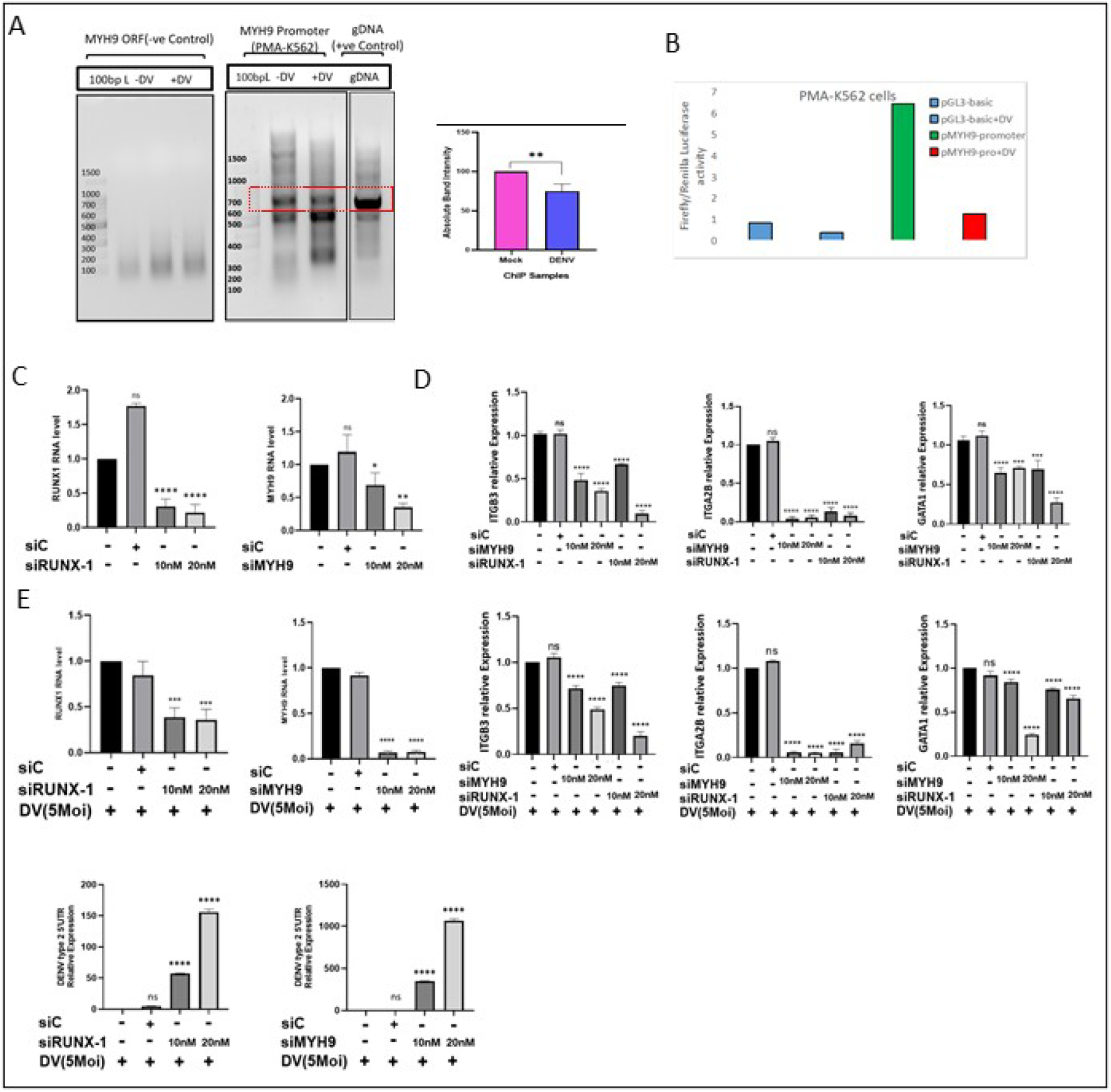
RUNX-1-mediated transcriptional regulation of MYH9 is perturbed during DENV infection in the K562-megakaryocytic model. RUNX-1-mediated MYH9 gene regulatory axis has been evaluated in a DENV-infected PMA-K562 cell line. (A) MYH9-promoter DNA was precipitated with RUNX-1 antibody in vitro from PMA-K562 cells, and the precipitated DNA was subjected to RT-PCR analysis. ChIP data suggest a decrease in the physiological association between the MYH9 promoter and the RUNX-1 transcription factor during dengue virus infection. (B) PMA-K562 cells were transfected with an empty pGL-3-basic vector as a vector control (VC) at a concentration of 1 µg or with a plasmid carrying MYH9-promoter (pMYH9-promoter) at a concentration of 1 µg. Twenty-four hours post-transfection, cells were infected with DENV at a 5 MOI, and cells were harvested 24 hours post-infection in Passive Lysis Buffer. The cells were subjected to a luciferase assay. The analysis indicates a decrease in MYH9 Promoter activity during DENV infection. (C) PMA-K562 cells were mock-transfected or transfected with non-specific siCtrl (siC), siRUNX-1, and siMYH9 at concentrations of 10nM and 20nM. Cells were harvested at 3 days post-transfection and were subjected to MYH9 and RUNX-1 expression analysis. qRT results indicate that RUNX-1 knockdown cells displayed a reduction in reduced RUNX-1 expression. Additionally, the MYH9 knockdown in siMYH9 cells was confirmed with real-time PCR. (D) Megakaryopoiesis-proliferative marker gene expression was also investigated in RUNX-1- and MYH9-knockdown cells. The qRT-PCR data indicate reduced megakaryocytic differentiation under knockdown conditions. (E) Similarly, megakaryopoiesis-proliferative marker gene expression has been assessed in RUNX-1- and MYH9-knockdown cells infected with dengue virus. The data indicate that megakaryopoietic gene expression is down-regulated under both conditions compared to control cells.

Furthermore, we have taken MYH9-ORF as a negative control. We performed RT-PCR with the MYH9-ORF region, and we did not observe amplification of the MYH9-ORF region from the precipitated samples, which suggests the specificity of RUNX-1 binding to the MYH9-promoter region. Additionally, genomic DNA (gDNA) amplification was taken as a positive control. Our observation indicates that DENV infection modulates the RUNX-1 interaction with the MYH9 promoter in both the Huh7 cell line and the K562 cell line. This leads to dysregulated platelet biogenesis.

After establishing through the ChIP Assay that the binding of RUNX-1 to the promoter region of the MYH9 gene is altered upon DENV infection. We next aim to investigate whether this alteration affects MYH9 promoter activity. To determine this, a luciferase reporter assay was implemented. The cloned MYH9 promoter in the pGL3-Basic vector was utilized to investigate the transcriptional activity of the MYH9 promoter under mock- and DENV-infected PMA-K562 cells to quantitatively measure the changes in MYH9 promoter activity. Our data indicate a comparable trend in PMA-Induced K562 cells. Under mock-treated conditions, robust promoter activity was detected. Whereas a significant decrease in the luciferase activity was observed (Figure 5B), further supporting the hypothesis that DENV-mediated perturbation of RUNX-1 binding compromises the transcriptional activity of the MYH9 promoter.

Subsequently, we sought to determine the regulatory effect of RUNX-1 knockdown on megakaryopoietic gene expression. For this, we knocked down RUNX-1 using siRNA specific against RUNX-1 in PMA-K562 cells. The K562 cells were seeded and transfected with siC and siRUNX-1 at final concentrations of 10nM and 20nM; the mock-transfected cells were taken as a control. After transfection, the cells were induced with PMA at a concentration of 50nM. The cells were harvested at 3 days post-induction, and RUNX-1 mRNA levels were assessed by quantitative PCR. Our observation indicates a concentration-dependent reduction in RUNX-1 mRNA levels under knockdown conditions (Figure 5C).

Since MYH9 is an important protein that functions in platelet biogenesis, we suspect that downregulation of MYH9, either in direct knockdown (Figure 5C) or in RUNX-1 knockdown cells, affects the megakaryopoiesis process. To this end, we further assessed the effects of RUNX-1 and MYH9 knockdown on the megakaryopoiesis-associated proliferative marker genes CD61, CD41, and GATA. siMYH9 mediated MYH9 downregulation has been performed in PMA-k562 cells. The samples were harvested at 3 days post-induction, and expression of the proliferative marker gene was quantified with real-time PCR. The data show a reduction in the expression of CD61, CD41, and GATA1 under both RUNX-1 and MYH9 knockdown conditions (Figure 5D). This observation suggests that MYH9 downregulation could significantly impair megakaryopoiesis.

Similarly, MYH9- and RUNX-1-knockdown cells were subjected to dengue infection at 5 MOI 24 hours post-transfection, followed by PMA induction. The samples were harvested and evaluated for megakaryopoitic proliferative marker gene expression. Quantitative real-time PCR shows reduced expression of megakaryopoiesis-related genes, indicating dysregulation of megakaryopoiesis during dengue virus infection (Figure 5E). In siMYH9 and siRUNX-1 cells, we examined dengue virus RNA levels; for this, we quantified viral RNA by qPCR using primers specific to the DENV type 2 5’UTR. The observation indicates elevated dengue viral RNA levels, suggesting that MYH9 and RUNX-1 down-regulation aids viral proliferation.

### MYH9 overexpression rescues cytoskeletal anomalies in the DENV-infected K562 Megakaryocytic model

Since K562 cells are suspension cells and inherently difficult to transfect, it was essential to first establish optimal transfection conditions. We focused on optimizing the transfection strategy for both uninduced and PMA-induced K562 cells (details are provided in Supplementary Figure S5).

Furthermore, the pV5-MYH9-WT plasmid construct (Addgene no.183512) was overexpressed in PMA-K562 cells using strategy 2. The recombinant plasmid was transfected at concentrations of 0.5 µg, 1 µg, and 2 µg, while cells transfected with an empty vector at a concentration of 2 µg were used as a transfection control (VC). The exogenous expression of MYH9 protein was confirmed by immunoblotting using an anti-V5-tag antibody (Figure 7A (i)) along with endogenous MYH9 detection with an anti-MYH9 antibody. The recombinant plasmid promoted transient expression of MYH9-WT under the control of the CMV promoter. Moreover, the expression of MYH9-WT was evaluated at 2 and 3 days post-PMA induction in K562 cells by immunoblotting. The results show sustained MYH9 overexpression up to 3 days post-PMA induction (Figure 6A(ii)).

**Figure 6.**
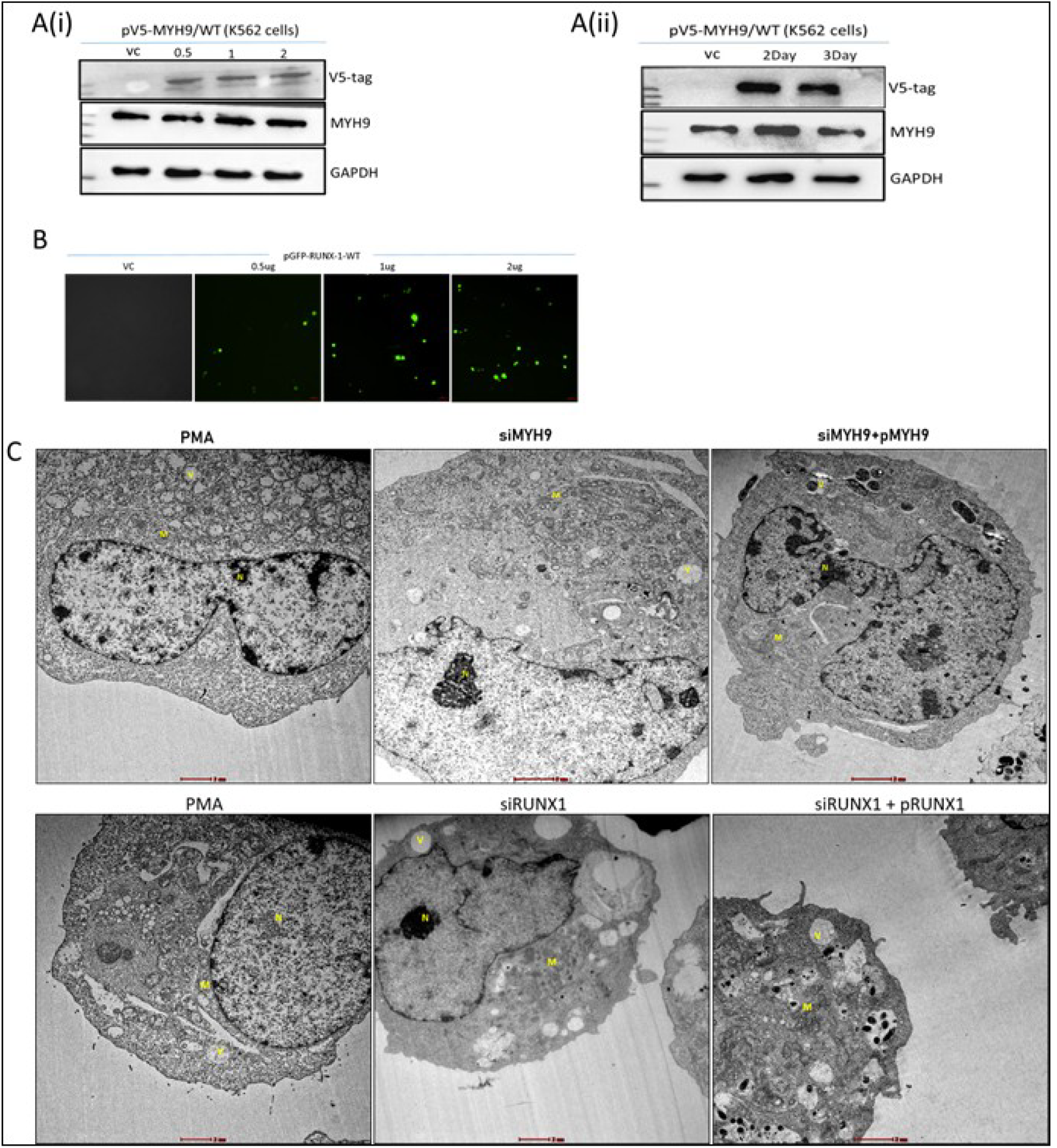
MYH9 overexpression rescues the cytoskeletal anomalies in the DENV-infected K562-megakaryocytic model. Ultrastructural analysis was performed on PMA-induced K562 cells for the rescue experiment. (A) pV5-MYH9-WT plasmid construct was transfected into PMA-induced K562 cells. The cells transfected with an empty vector were taken as a vector control (VC). (i) The cells were transfected with pV5-MYH9-WT at concentrations of 0.5 µg, 1 µg, and 2 µg. The overexpression of MYH9 protein was confirmed with Western blotting using a V5-tag antibody. (ii) Additionally, MYH9 overexpression was assessed at 2-day and 3-day post-induction. GAPDH was taken as a loading control. (B) The pGFP-RUNX1-WT plasmid construct was transfected into PMA-induced K562 cells. The cells transfected with an empty vector were taken as a vector control. RUNX1 gene expression was confirmed by the detection of GFP at 3 days post-PMA induction. (C) TEM-based ultrastructural analysis was performed from PMA-only, MYH9, and RUNX1 knockdown cells. The analysis indicates partial restoration of differentiated megakaryocytic features in KD cells overexpressing MYH9 and/or RUNX1. The markings N, V, and M represent the nucleus, vacuole, and mitochondria, respectively.

**Figure 7.**
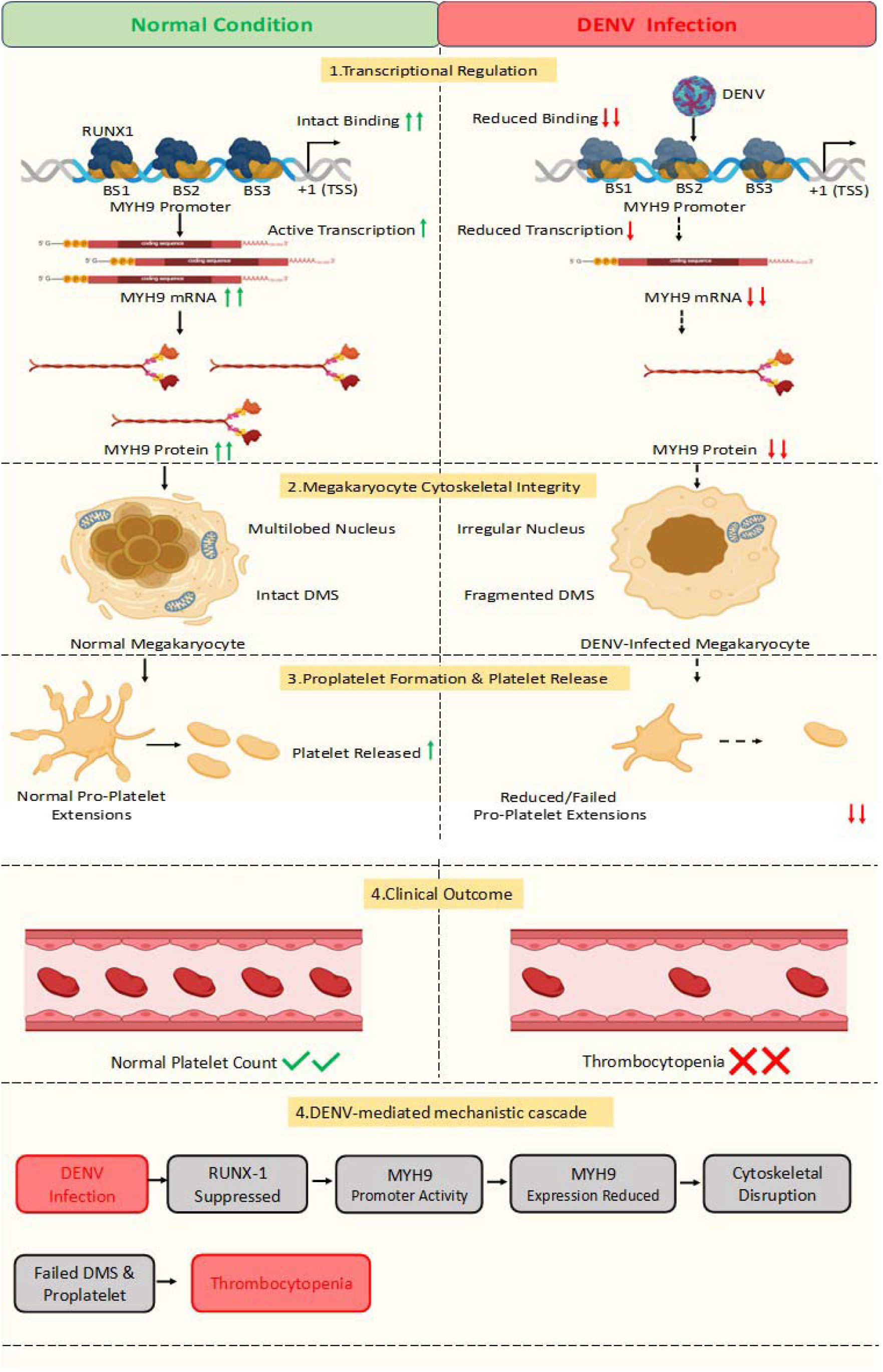
Summary. This illustration depicts that upon DENV infection, the interaction between RUNX-1 TF and the MYH9 promoter gets disrupted, leading to perturbation in the functional as well as expressional regulation of the MYH9 gene, resulting in a reduction in the generation of MYH9 transcripts, thereby causing a reduced production of MYH9 protein, which ultimately leads to the disruption of cytoskeletal integrity of platelets, which is a major contributor to thrombocytopenia. Abbreviations: BS: Binding Site, TSS: Transcription Start Site, DMS: Demarcation Membrane System

Moreover, the pGFP-RUNX1-WT plasmid construct (Addgene no.205915) was overexpressed in PMA-K562 cells using strategy 2. The recombinant plasmid was transfected at concentrations of 0.5 µg, 1 µg, and 2 µg, while cells transfected with an empty vector at a concentration of 2 µg were used as a transfection control (VC). The exogenous expression of RUNX1 protein was confirmed by immunofluorescence by observing the GFP expression (Figure 6B).

Subsequently, to examine the functional reversal of MYH9 and RUNX1 proteins in siMYH9 and siRUNX1 cells, we evaluated ultrastructural features using transmission electron microscopy (TEM) in MYH9- and RUNX1-overexpressing cells. Briefly, K562 cells were seeded at 100 X 10^4^ cell density and were transfected with siMYH9 or siRUNX1 at 20nM concentration. Twelve hours post-transfection, the respective KD cells were transfected with p-V5-MYH9 or p-GFP-RUNX1 plasmid constructs at a concentration of 1 μg. Then, 7-8 hours post-transfection, the cells were treated with 50 μM PMA and incubated for 3 days before proceeding with the TEM protocol. PMA-alone-treated cells, MYH9 KD cells, and RUNX1 KD cells were used as controls for comparison of ultrastructural features.

Primarily, our TEM analysis demonstrates that PMA-treated K562 cells exhibit hallmark features of megakaryocytic differentiation, including well-defined cytoplasm, increased mitochondrial biogenesis, vacuolization, cellular membrane trafficking, and the formation of an extensive plasma membrane-derived demarcation membrane system (DMS). The ultrastructural analysis of MYH9 knock-down cells reveals unstable cytoplasmic architecture, decreased vesicle biogenesis, mitochondrial accumulation due to reduced cellular organelle trafficking, and fragmented, discontinuous DMS formation, as shown in Figure 6C. This mechanical instability has been anticipated as a result of reduced MYH9 protein function; MYH9 encodes non-muscle myosin IIA, a key motor protein responsible for actomyosin contractility, intracellular tension, and membrane organization. Loss of MYH9 therefore compromises the mechanical framework required to transform early differentiation signals into stable ultrastructural architecture. TEM analysis of the siMYH9 K562 cells overexpressing MYH9 protein shows partial functional restoration of MYH9 protein. Our TEM analysis demonstrated a more uniform cytoplasm with better organelle distribution and cytoplasmic organization, which is closer to normal differentiated morphology. Another key differentiating feature observed is a bilobed nucleus, as shown in Figure 6C (upper panel).

In parallel, we have also explicated the inevitable role of RUNX1 transcription factor in megakaryocytic differentiation. We have compared the ultrastructural morphology of siRUNX1 KD-only cells with RUNX1 protein overexpressing siRUNX1 KD cells. Our observation indicates that RUNX1 knockdown cells typically initiate megakaryocytic differentiation but fail to undergo megakaryocytic maturation. PMA induction initiated cellular expansion and vesicle formation, which are early differentiation signals, but also impaired the formation of the internal membrane system (DMS), leading to irregular nuclear boundaries, abnormal organelle organization, and larger vacuole formation, as shown in Figure 6C. Our rescued cells exhibited a membrane network system, granule formation, better-organized mitochondrial distribution, and structurally organized cytoplasm. These observed features suggest a differentiating state in RUNX1-overexpressing cells, as shown in Figure 6C (lower panel).

Altogether, our TEM ultrastructural analysis revealed restoration of the megakaryocytic differentiation state compared with typical MYH9 or RUNX1 knock-down features.

## DISCUSSION

Dengue is a vector-borne disease that is persistent worldwide. Dengue infection causes dengue fever or more life-threatening dengue shock syndrome or dengue hemorrhagic fever (31). Severe DENV infection manifests with a clinical hallmark condition, that is thrombocytopenia, which is identified as a major drop in platelet counts contributing to the severity and making thrombocytopenia a major health concern for patients, yet the molecular mechanisms underlying this condition remain to be elucidated (32).

Previously, one of the studies established the genetic mutations in 41 genes from the platelet biogenesis pathway linked to inherited bleeding disorders and defective megakaryopoisis differentioation/maturation (33). With this foundational literature, we sought to document the link between these megakaryopoitic genes and dengue-associated thrombocytopenia. Primarily, we performed expression profiling of a subset of these documented genes in a Huh7 dengue-permissive cell line. Gene expression profiling of megakaryopoiesis revealed significantly reduced MYH9 expression during dengue virus infection at both the RNA and protein levels.

MYH9, a non-muscle myosin heavy chain II-A protein. MYH9 participates in maintaining cellular cytoskeletal integrity, cell motility, and cell shape (34). Interestingly, platelets exclusively produce the MYH9 isoform. This MYH9 protein is critically required for megakaryocytic maturation and proplatelet formation (35). Therefore, we hypothesized that any change in the expression or cellular localization of this protein would severely affect platelet biogenesis (Figure 7). Previous studies have documented the role of the MYH9 protein in various viral infections, including HSV-1 (36), PRRSV (37), Influenza A virus (29), SARS-CoV-2 (38), and Dengue virus (39). These studies documented the pro-viral activity of MYH9 by acting as a membrane receptor to facilitate viral attachment/entry during viral infection, establishing MYH9 as a broad-spectrum viral entry agent (28). The study on dengue virus also suggests the pro-viral activity of MYH9; it has shown increased expression of MYH9 and its relocalization to the cell surface, where it acts as a receptor for dengue virus internalization (39). Recently, a study by Chen et al. has shown the dual regulatory function of the MYH9 protein in IAV infection. They have demonstrated that MYH9 accumulates at the cell surface to facilitate viral attachment and also inhibits viral replication by interfering with the viral replication complex; thus, it can act as both a pro-viral and an anti-viral factor (29). However, none of the studies have elucidated MYH9 expression and its regulation. On the other hand, previous studies have demonstrated an association between MYH9 depletion and inherited thrombocytopenia or bleeding disorders (40,41). For instance, inherited bleeding disorders including May-Hegglin anomaly (MHA), Sebastian syndrome (SBS), Fetchtner syndrome (FS), and Epstein syndrome (ES) have been shown to result from MYH9 mutations, leading to decreased or absent MYH9 expression/function, leading to macrothrombocytopenia, ultrastructurally defective megakaryocytes, early release of immature platelets, and terminal defects in platelet biogenesis (42). These disease conditions are collectively referred to as MYH9-related disorders (MYH9-RD). In this study, we elucidated the intricate transcriptional regulation of the MYH9 gene in dengue-associated thrombocytopenia.

Subsequently, RUNX-1 has been identified as a transcriptional activator of the MYH9 and MYL9 genes, which are required for cytoskeletal assembly in platelets. It is a master gene regulatory agent critical for megakaryocyte lineage commitment (43). Our *in-silico* analysis from the eukaryotic promoter database has revealed three potential binding sites for the RUNX-1 transcription factor in the MYH9 promoter region. We further demonstrated the physiological association between RUNX-1 and the MYH9 promoter, as well as promoter activity. The dengue virus modulates the interaction between RUNX-1 and the MYH9 promoter, thereby reducing MYH9 promoter activity. The knockdown studies further revealed that the MYH9 gene is indeed under the control of the RUNX-1 transcription factor. We have demonstrated the concomitant reduction of MYH9 gene expression with decreased RUNX-1 expression in a concentration-dependent manner. The dengue virus further substantiates this effect.

MYH9 is a cytoskeletal protein, and hence the depletion of the MYH9 protein has been shown to disrupt the cytoskeletal organization and cell migration (44,45). The immunofluorescence images in our data also show cytoskeletal organization defects, with a compromised cytoskeletal appearance and disorganized actin fibers more concentrated towards the periphery rather than distributed throughout the cells. This observation suggests that decreased MYH9 expression affects cytoskeletal organization, leading to cytoskeletal anomalies that are required to maintain cellular shape and motility.

Given that Huh7 cells are non-megakaryocytic cell lines and we sought to determine the effect of decreased MYH9 on megakaryocyte maturation, we used K562 cells, which can mimic megakaryocytic differentiation in vitro.

Recently, an RNA-seq study of DENV-infected K562 cells revealed that dengue virus infection upregulates the NFE2L2 gene to suppress ROS accumulation and promote its own replication (18). Since the study uses a PMA-induced K562 cell line model that recapitulates megakaryocytic differentiation and maturation, we reanalyzed the sequencing data to assess differential expression of megakaryopoietic genes. The heatmap shows that the expression of many megakaryopoietic genes is altered during dengue virus infection. Moreover, we validated the RNA-seq results using the proliferative marker genes GATA1, ITGA2B (CD41), ITGB3 (CD61), the cytoskeletal marker gene MYH9 (our leading candidate), and the apoptotic marker gene CYCS in K562-megakaryocytic cells. Our data indicate upregulation of PMA differentiation-associated proliferative and cytoskeletal genes, while apoptotic gene expression decreases during megakaryocytic maturation. Conversely, during DENV infection, the expression of proliferative and cytoskeletal marker genes is reduced, and the apoptotic gene CYCS is upregulated, suggesting a multi-channel effect of dengue virus on platelet biogenesis. The virus selectively downregulates the pro-platelet factor while increasing the levels of the apoptotic gene CYCS to augment platelet clearance. Additionally, simple microscopic observation of K562 cells, PMA-K562 cells, and dengue-infected PMA-K562 cells clearly reveals altered megakaryocytic differentiation dynamics. The only K562 cells (control cells) are visibly smooth, round, small, and non-adherent; the PMA-induced K562 cells exhibit increased cell size and surface adherence, with a vacuolated morphology. Interestingly, dengue-infected PMA-K562 cells exhibit altered morphology and reduced adherence, indicating impaired megakaryocytic differentiation.

Moreover, we established a RUNX-1-mediated regulatory axis of MYH9 gene expression in K562-megakaryocytic cells. The megakaryopoietic gene expression profiling revealed significantly reduced MYH9 expression during dengue virus infection. Similarly, we established RUNX-1-mediated regulation of the MYH9 gene and the modulatory effect of dengue virus on this regulatory axis. The data in this report show that MYH9 expression is under the transcriptional control of RUNX-1 in megakaryocytic cells, and that DENV infection modulates this regulatory axis to dampen MYH9 expression. Quantitative PCR data from RUNX-1 and MYH9 knockdown studies show reduced expression of megakaryopoietic genes, including CD61, CD41, and GATA1. This observation suggests the crucial roles of MYH9 and RUNX1 in megakaryocytic differentiation, and that their reduced expression leads to aberrant megakaryocytic differentiation signals. Altogether, this report indicates that dengue virus modulates MYH9 function in megakaryocytic cells as well as non-megakaryocytic permissive cells.

Importantly, in this report, ultrastructural images of RUNX-1- and MYH9-knockdown cells revealed reduced megakaryocytic differentiation and maturation signals compared with PMA cells. Specifically, siMYH9 cells exhibit disorganized cellular trafficking due to structural defects, whereas siRUNX-1 cells show lineage-commitment defects. The PMA cells show the typical PMA differentiation features, including bilobed nucleus, cellular organelle distribution, and demarcation membrane formation. In addition, the overexpressing cells partially restore PMA differentiation and maturation signals. This provides evidence that the dengue-associated decrease in platelet count may be restored, thereby partially reversing dengue-associated pathogenic effects.

Overall, our findings provide compelling evidence that DENV infection impairs the process of platelet biogenesis by perturbing the RUNX-1-MYH9 regulatory axis. This perturbation compromises cytoskeletal integrity and impairs megakaryocyte maturation and proplatelet formation, ultimately leading to thrombocytopenia. By shedding light on this molecular mechanism, our study highlights potential therapeutic strategies for mitigating platelet depletion during dengue infection.

## Acknowledgment

The authors acknowledge All India Institute of Medical Sciences, New Delhi, and Department of Biotechnology, AIIMS, New Delhi for the support.

## Author contributions

A.S. and B.V.: conceptualization, methodology, and formal analysis; A.S., S.M., K.S., S.P., A.K., V.C., S.B., and B.V.: investigation; A.S., S.M., K.S., and B.V.: writing– original draft; B.V.: supervision; B.V.: funding acquisition.

## Data availability

All data used to support the findings of this study are included within the article.

## Supplementary Information

Supplementary figures 1-5

