## Supplementary Data for "Dengue virus-induced thrombocytopenia is mediated by dysregulation of the RUNX1–MYH9 axis"

1. **MYH9 cloning Method:**

**Supplementary figure S1**

**MYH9-promoter cloned between – KpnI and HindIII**

**Primers:**

**5’-TATTGGTACCGAACGGCCAGAGCGAAGGTTTG-3’**

**5’-TATTAAGCTTGCGCGCCTCACCGCACCCA-3’**

**Reaction condition:**

1) **Insert Amplification from genomic DNA.**

Initial Denaturation: 98°C- 3min

Denaturation: 98°C- 30sec

Annealing: 58.8°C- 20sec

Extension: 72°C- 30sec

Final Extension: 72°C- 5min

Cooling: 4°C

2) **Vector and insert double digestion reaction.**

(Both enzymes are from NEB)

Buffer: 2ul

Enz. kpnI: 1ul

Vector DD

Enz. HindIII: 1ul

(800ug) pGL-3-basic: 3.2ul

NFW: 12.8ul

Buffer: 2ul

Enz kpnI: 1ul

Insert DD

Enz HindIII: 1ul

(400ng) MYH9-Pro: 12ul

NFW: 4ul

Digestion Reaction has been incubated overnight at 37°C for 16hrs followed by enzyme inactivation at 65°C for 20 mins.

3) **Vector and insert ligation reaction.**

10X T4 DNA ligase Buffer: 2ul

T4 DNA ligase enz: 1ul

DD vector: 1ul

Insert (5:1):

NFW:

Nanodrop reading of DD vector= 35ng/ul

Nanodrop reading of DD insert= 10.4ng/ul

Insert volume calculation for 5:1 ligation :-

i) Using NEB calculator= Required insert mass is 25.48ng
ii) Manual calculation (V mass X 5)/insert mass = 16ul.

Number of reaction:-

1 V+I

2 V+I

3 V- I

4 UD Vector

Ligation reaction has been incubated O/N at 16°C for 16hrs followed by enzyme inactivation at 65°C for 20mins.

1. MYH9 promotor amplification using MYH9 (pro) primers

From genomic DNA isolated from cell line sample

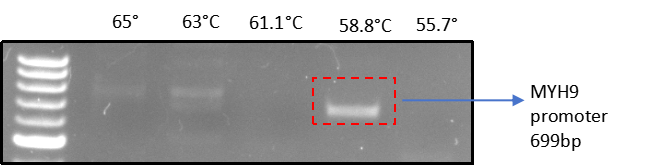

2. Double digestion of vector and insert

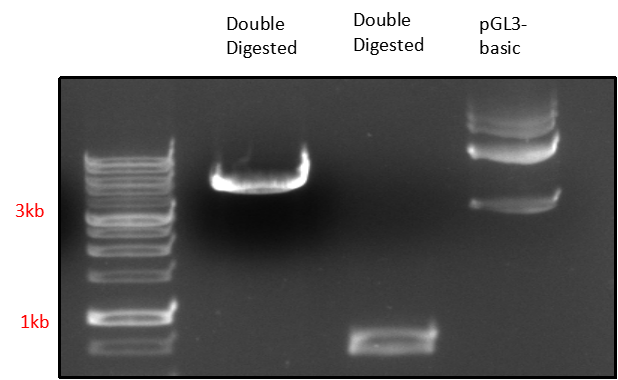

3. Double digestion screening of Pgl3-basic-MYH9pro construct

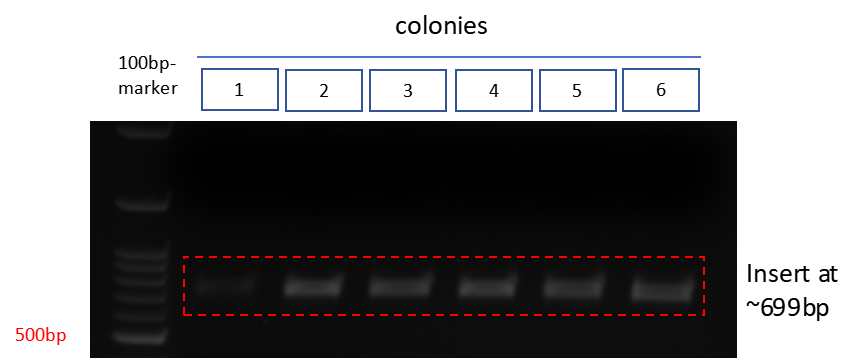

4. PCR screening of Pgl3-basic-MYH9pro construct

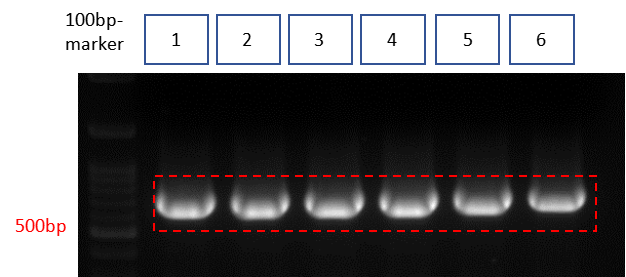

**Sanger Sequencing Result:**

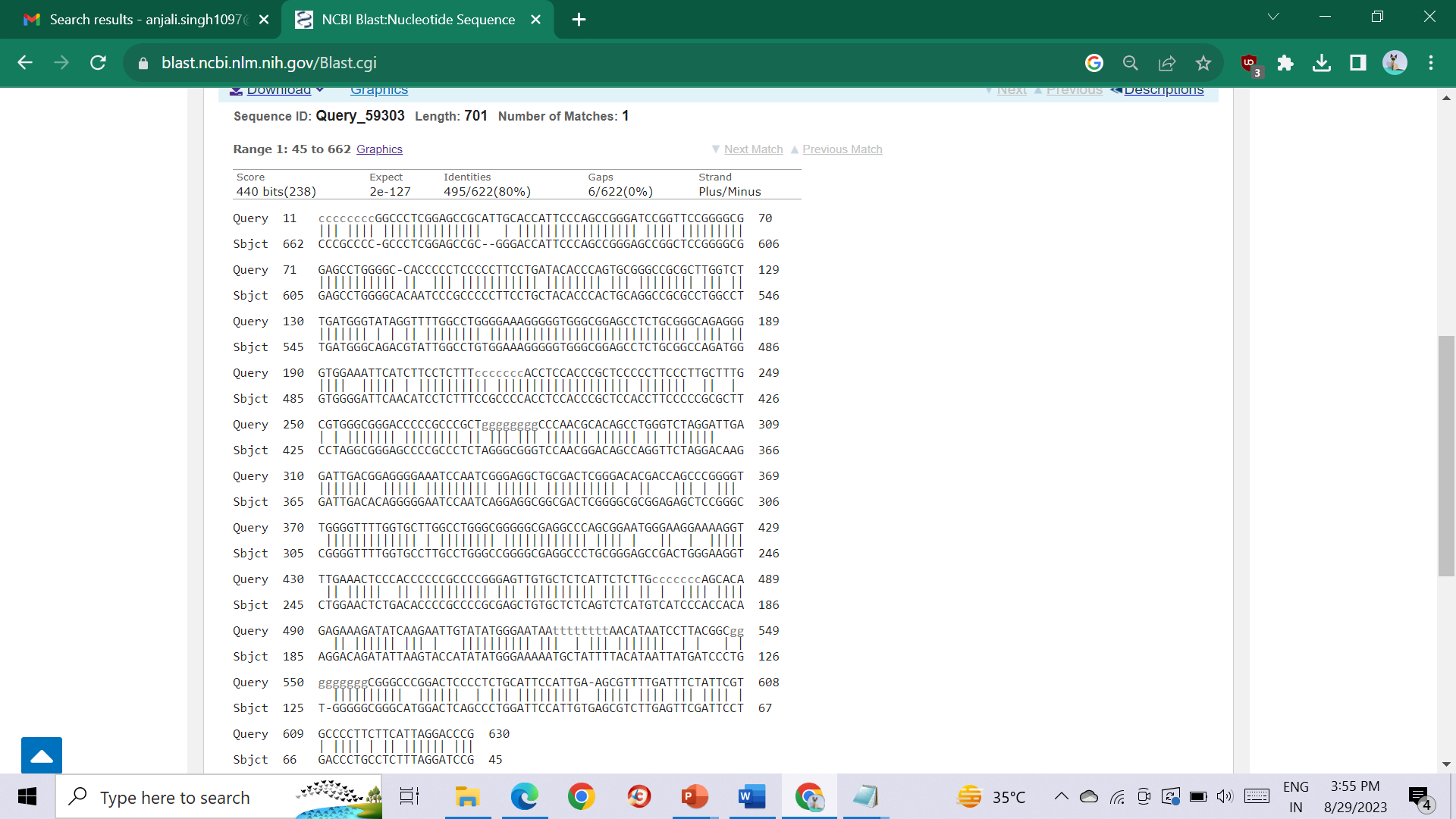

**Supplementary Figure S1: MYH9-promoter cloning strategy and Sanger sequencing output.**

2. **Viral infection detection using qRT-PCR:**

**Supplementary figure S2**

The dengue virus infection has been confirmed in dengue MOI samples using primers specific against dengue envelop region**.** The PCR analysis indicates MOI dependent enhancement in viral RNA detection.

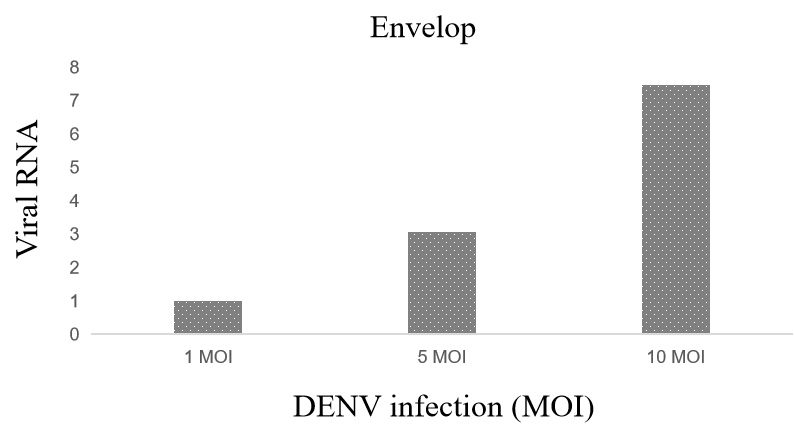

**Supplementary figure S2**: Viral RNA detection in dengeu inefcted Huh7 cells.

3. **Sample grouping for the RNA-seq data:**

**Supplementary figure S3**

To examine the differential expression of Megakaryopoitic genes in PMA-induced K562 cells, we re-analyze the RNA-seq data submitted to GEO Database (GSE186089) from Dr. Sankar Bhattacharyya’s Lab (THSTI, Faridabad, India). they have taken total 12 RNA samples for RNA Sequencing: A- 0 Day PMA Induced, Mock infected (in duplicates); B- 3 Day PMA Induced, Mock infected (in duplicates); C-3 Day PMA Induced, DENV infected (in triplicates), D- 6 Day PMA Induced, Mock infected (in duplicates); D- 6 Day PMA Induced, DENV infected (in triplicates).

For the purpose of RNA-seq analysis, samples were grouped into certain sets for relevant comparison. Total 4 sets were formed for the analysis as depicted in figure 6 B(i). **Set-A B**, to indicate the changes observed in uninfected cells after 3 days of PMA-induced differentiation (set B) when compared to 0-day samples (set A). **Set-B C**, to indicate the changes observed between uninfected (set B) and DENV-infected cells (set C) after 3 days PMA-induced differentiation. **Set- A D**, indicates the changes observed in uninfected cells after 6 days of PMA-induced differentiation (set D) when compared to 0-day samples (set A). **Set-E D**, to indicate the changes observed between uninfected (set D) and DENV-infected cells (set E) after 6-days PMA-induced differentiation.

Following this, differential gene expression analysis was performed to identify genes and pathways altered across experimental conditions. Raw RNA sequencing reads were aligned to the most recent stable version of the human reference genome GH38 (GRCh38.p5, Ensembl) using Bowtie 2 and Tophat 2.1.1. Count data was normalized across samples and differential expression analysis conducted using the R package DESeq2. For each sample, normalized gene and transcript expression profiles were computed. The FPKM (Fragments Per Kilobases per Million fragments) method was used followed by e log2fold change transformation of the value. The gene-level differential expression in different conditions were estimated using the log2 transformed FPKM. While identifying differentially expressed genes (DEGs), uncorrected p-value of the test statistic and the FDR-adjusted p-value of the test statistic (q-value) were also calculated. After Benjamini-Hochberg correction for multiple testing, any gene with a p-value higher than the FDR was considered significantly differentially expressed.

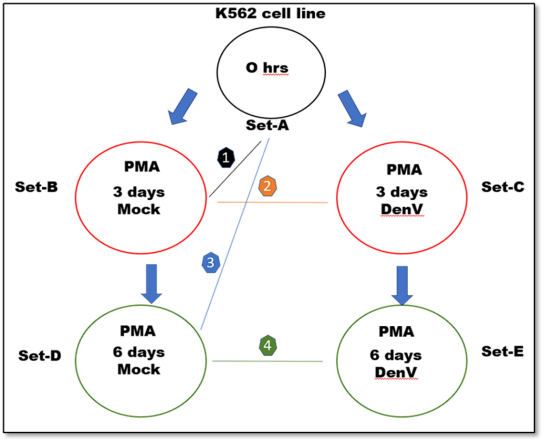

**Supplementary Figure S3: Sample grouping of RNA-seq analysis.**

4. **Sodium butyrate control PMA induced K562 model establisment:**

**Supplementary figure S4**

In addition to PMA induction and DMSO vehicle control, a parallel set of K562 cell cultures were treated with 20 mM sodium butyrate over the 6-day time course. As a histone deacetylase inhibitor, butyrate elicits gene expression changes in K562 cells, notably stimulating erythroid differentiation markers. By propidium iodide polyploidy profiling, 0 hr butyrate-treated cells matched DMSO and PMA baseline profiles with ~61% 2N fraction indicating a progenitor state. At day 3 post-treatment and even further, majority of the populations exhibited a ploidy of 2N, with no considerable 4N or higher peaks observed as compared to PMA induced K562 cell samples.

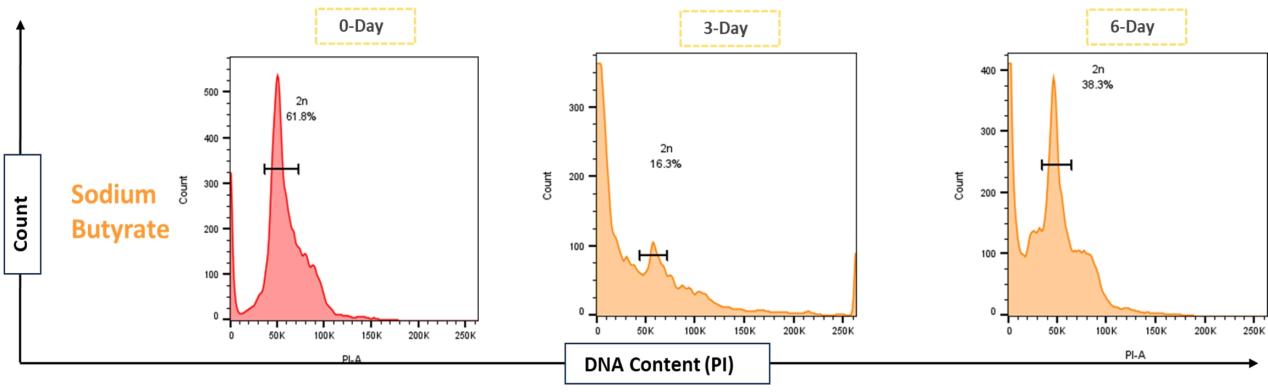

**Supplementary Figure S4: Flow cytometry using PI staining of sodium-butyrate induced K562- megakaryocytic cells do not show any changes in ploidy of the megakaryopoitic cells.**

5. **Transfection strategy to transfect plasmid construct in K562 cells:**

**Supplementary figure S5**

To achieve transfection in suspension cells, we systematically tested and refined various transfection strategies to enhance efficiency and reprehensibility before proceeding with further functional assays.

To optimize transfection conditions in uninduced K562 cells, two different transfection reagents, LTX and Lipofectamine 2000 (LPF), were evaluated using a GFP-tagged plasmid. Immunofluorescence microscopy analysis performed at 24- and 48-hours post-transfection revealed a higher GFP signal in LPF-transfected cells compared to those transfected with LTX, indicating superior transfection efficiency. Quantitative analysis further confirmed this trend, demonstrating significantly enhanced transfection efficiency with LPF. These results establish Lipofectamine 2000 as the more effective reagent for transfecting K562 suspension cells, making it the preferred choice for subsequent experiments.

To determine the optimal transfection strategy for PMA-induced K562 cells, two different approaches were tested using Lipofectamine 2000 (LPF) as shown in figure. Strategy 1: K562 cells were first treated with PMA for 2 hours, after which the PMA was removed, and transfection with LPF was performed. Cells were imaged on the 3rd day post-transfection. Strategy 2: K562 cells were transfected with LPF first, followed by PMA induction 8 hours post-transfection. Imaging was captured on the third day post-transfection. Both strategies achieved good transfection efficiency. However, we proceeded with Strategy 2 as it resulted in a more robust and well-established PMA-induced differentiation, making it the preferred method for further experiments.

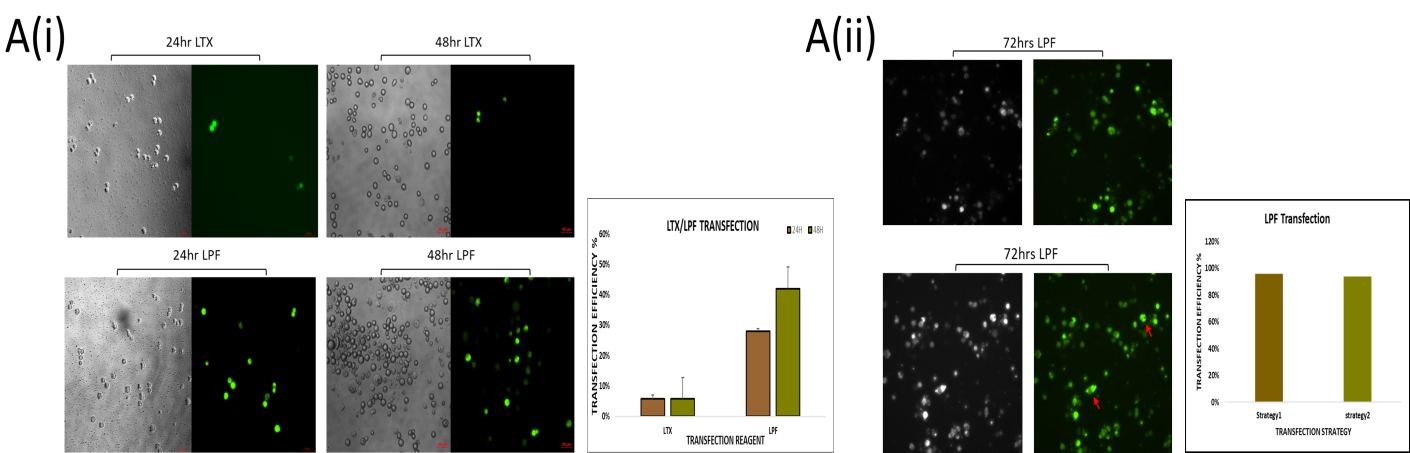

**Supplementary Figure S5: Transfection strategy was optimized in PMA induced K562 cells for overexpression studies.** (A)(i) K562 cells were seeded in 96wells plate at 3 X 10^4^ cell density followed by transfection with LTX and Lipo-2000 in separate wells with the pGFP-RUNX-1 plasmid construct. The cells were observed under immunofluorescence microscope at 24 hpt and 48 hpt to visualize transfection efficiency. The result shows that Lipo-2000 transfected cells were showing higher GFP expression compared to LTX transfecting Reagent. (ii) K562 cells were seeded into 96 wells plate followed by Lipo-2000 mediated transfection of pGFP-RUNX-1. The 8 hours post transfection, cells were induced with 50nM PMA. The cells were then observed at 72 hours post induction for transfection efficiency.

6. **Primers Sequences:**

**Supplementary table 1**

| Genes | Primer pair | Primer sequence |
| --- | --- | --- |
| GATA-1 | GATA-1 F | 5’-CACTACCTATGCAACGCCTG-3’ |
|  | GATA-1 R | 5’-TGTAGCTTGTAGTAGAGGCCG-3’ |
| ETV6 | ETV6 F | 5'-TGTAGCATTAAGCAGGAACGAA-3 |
|  | ETV6 R | 5'-TCCTGGGACTCTAGGTGCTC-3' |
| RBM8A | RBM8A F | 5'-ATGGCGGACGTG CTAGAT-3' |
|  | RBM8A R | 5'-CTGGACTTCTGCTGCGTCTT-3' |
| ACTNl | ACTNl F | 5'-TGGACCATTATGATTCTCAGCA-3' |
|  | ACTNl R | 5'-TTAGAGGTCACTCTCGCCGTA-3' |
| TRPM7 | TRPM7 F | 5'-TGTCCCAGAAATCCTGGATAG-3' |
|  | TRPM7 R | 5'-TAACATCAGACGAACAGAATTAGTTG-3' |
| ABCG8 | ABCG8 F | 5'-TGGGTGACCTCTCATCTTTG-3' |
|  | ABCG8 R | 5'-TGCTAATGAGATGATCCCTTATTTT -3' |
| RUNXl | RUNXl F | 5'-GGCTTCAGACAGCATATTTGAG-3' |
|  | RUNXl R | 5'-CAGTAGGGCCTCCACACG-3' |
| ITGB3 | ITGB3 F | 5’-AGTGAGGCCCGAGTACTAGA-3’ |
|  | ITGB3 R | 5’-AGTTACTGGTGAGCTTTCGC-3’ |
| CYCS | CYCS F | 5’-ATGGTGATGATGTTGAGAA-3’ |
|  | CYCS R | 5’-TAAGTCTGCCCTTTCTTCC-3’ |
| ITGA2B | ITGA2B F | 5’-TGCTCTTTGACCTCCGTGAT3-3’ |
|  | ITGA2B R | 5’-CGCTTCACAGTAACGCTTGT-3’ |
| MYH9 | MYH9 F | 5’-CAAGAAGCTGGTATGGGT-3’ |
|  | MYH9 R | 5’-TGCCTCTTCTTGCCCTTGT-3’ |

**Table 1:** List of primer sequences of the megakaryopoitic genes.
